# Stereochemistry of transient protein-protein interactions in a signaling hub: exploring G3BP1-mediated regulation of CFTR deubiquitination

**DOI:** 10.1101/2022.03.02.482667

**Authors:** Colin T. Sheehan, Dean R. Madden

## Abstract

Intrinsically disordered proteins (IDPs) can coordinate often transient or weak interactions with multiple proteins to mediate complex signals within large, reversible protein networks. Among these, the IDP hub protein G3BP1 forms protein complexes with Caprin1 and USP10, and the resulting control of USP10 activity plays an important role in a pathogenic virulence system that targets CFTR endocytic recycling. However, while the identities of protein interactors are known for many of these IDP hub proteins, the relationship between pairwise affinities and the extent of protein recruitment and activity is not well understood. Here we describe *in vitro* analysis of the G3BP1 affinities, and show that substitution of G3BP1 residues F15 or F33 to tryptophan reduces affinity for both the USP10 and Caprin1 motif peptides. These same mutations significantly reduce formation of complexes by the full-length proteins. The crystal structure of G3BP1 TripleW (F15W/F33W/F124W) mutant reveals a clear reorientation of the side chain of W33, creating a steric clash with the USP10 and Caprin1 peptides. An amino-acid scan of the USP10 and Caprin1 motif peptides reveals similarities and differences in the ability to substitute residues in the core motifs as well as specific mutations with the potential to create higher affinity peptides. Taken together, these data show that small changes in 1:1 binding affinity can have significant effects on the composition of cellular interaction hubs. These specific protein mutations can be harnessed to manipulate complex protein networks, informing future investigations into roles of these networks in cellular processes.

## Introduction

Intrinsically disordered proteins (IDPs) are important molecular machines that play critical roles in cellular processes and signaling pathways (1, 2). Such proteins lack a well-defined three-dimensional globular structure and can range from fully to partially unstructured. This stereochemical flexibility allows these proteins to interact promiscuously with different proteins in a context-dependent manner (3). Additionally, these proteins often utilize conserved sequence motifs to facilitate protein-protein interactions with high specificity and modest affinity leading to rapid signal transduction (4–6). IDPs often function as interaction hubs as well as facilitators of biomolecular condensation. Ras GTPase Activating Protein SH3 Domain Binding Protein 1 (G3BP1), a protein containing many intrinsically disordered regions, is reported to serve as an interaction hub in stress granules (SG) and to form transient interactions with USP10 and Caprin1, which in turn influence the SG network organization and signaling (7–9).

G3BP1 is a 52 kDa protein ubiquitously expressed in the cytoplasm. Current research supports the theory that the main role of G3BP1 is to triage mRNA in response to intracellular and extracellular stimuli. However, significant evidence suggests that G3BP1 plays a central role in several additional cellular processes, including rasGAP signaling, ubiquitination, mRNA metabolism, and stress granule (SG) formation, and these diverse functions appear to be modulated by G3BP1’s interaction with RNA and other proteins (10–18). The large protein network surrounding G3BP1 has been implicated in several diseases including neurological and neurodegenerative disorders, cancer progression, bacterial pathogenesis, and viral infection (14, 16, 17, 19, 20). The ability to selectively modulate protein-protein interactions within this large, intricate protein network would represent a novel drug target with far reaching implications (21–24).

G3BP1 and the other two members of the G3BP family all share the following five domains: an N-terminal nuclear transport factor 2-like domain (NTF2), a central region consisting of several proline rich (PxxP) motifs, an acid-rich domain, a canonical RNA recognition motif (RRM), and a loosely conserved C-terminal arginine and glycine rich (RGG) box (25). Except for the NTF2-like domain and RNA-binding domain, G3BP proteins are predicted to be largely disordered. A recently reported model suggests that interactions with RNA and proteins partners modulate the structure and provide relative stability to G3BP1 (8, 9).

The NTF2 domain is most highly conserved domain in the G3BP family and plays a role in several G3BP functions including dimerization, protein binding, and SG formation (13, 26, 27). Proteins known to interact with G3BP1 NTF2 include rasGAP, Caprin1, and USP10 (10, 15, 28). Multiple groups have shown that Caprin1 and USP10 compete for the same binding groove on the G3BP1 NTF2 domain (7, 8, 29). Recent G3BP1 NTF2 co-crystal structures have identified three phenylalanine residues (F15, F33, and F124) that are responsible for coordinating peptides from Caprin1 and USP10 (26, 30). Additionally, several groups have shown that mutating F33 to a tryptophan ablates Caprin1 and USP10 binding (8, 26, 31, 32).

Ubiquitin Specific Peptidase 10 (USP10) is an 87 kDa cysteine protease and a member of the deubiquitinating enzyme (DUB) family. As a DUB family member, USP10 plays an important role in protein homeostasis and has been shown to remove conjugated ubiquitin from targets like p53/TP53, BECN1, SNX3 and the cystic fibrosis transmembrane conductance regulator (CFTR) (33–36). Additionally, USP10 has been implicated in a variety of diseases including cancer, where it can act as an oncogene or a tumor suppressor, as well as Alzheimer’s disease and other neurodegenerative diseases (19, 37–40).

USP10 contains a highly conserved catalytic domain (USP domain) beginning at residue R415 and spanning most of the C-terminal end of the protein. Similar to G3BP, the N-terminal half of the protein is predicted to be mostly disordered. First identified as a G3BP-interacting protein in a yeast two-hybrid screen, subsequent reports have identified a core motif (FGDF; residues 10-13) in USP10 that is recognized by G3BP1 NTF2 (15, 29). A co-crystal structure of NTF2 complexed with FGDF-containing peptides revealed that both motif phenylalanine residues (F10 and F13) protrude into the NTF2 binding grove and form π-stacking interactions with NTF2 residues F15, F33, and F124 (26, 32). Alanine substitution of USP10 F10, G11, or F13 completely ablates G3BP1 binding, while D12A allows for weak binding (32). These data show that the FGDF is necessary and sufficient for G3BP1 binding.

Cytoplasmic Activating/Proliferation Associated Protein 1 (Caprin1) is a 78 kDa cytoplasmic RNA-binding protein. Caprin1 has been implicated in cell-cycle regulation and cell proliferation where it acts alone or in combination with other RNA-binding proteins such as G3BP1 and fragile X mental retardation protein (FMRP) (28, 41–44). Caprin1 binds RNA via its C-terminal RNA-binding RGG motif and an RG enrichment region (28). Caprin1 contains a highly conserved F(M/I/L)Q(D/E)Sz(I/L)D motif spanning residues 372-379 that is recognized by G3BP1 NTF2 (28). A recent co-crystal structure revealed that the Caprin1 YNFI(Q) segment binds in the same hydrophobic grove on NTF2 as the USP10 FGDF motif (30). The structure further highlighted the important role of NTF2 residues F15, F33, and F124 in coordinating Caprin1 and USP10 peptides.

G3BP1 and its interactions with USP10 and Caprin1 are important for SG formation and regulation. Additionally, USP10 and Caprin1 have distinct roles in several cellular processes separate from G3BP1 and SGs (28, 33, 34, 45, 46). However, there is limited information on the importance of specific conserved motifs for the stability of IDP interactions. We therefore sought to better understand the interaction of these proteins at the residue level and to generate better tools for future investigations. We report here the first crystal structure of G3BP1 NTF2 harboring three phenylalanine substitutions in residues F15, F33, and F124 (TripleW). Additionally, we show a full amino-acid scan of the USP10 and Caprin1 binding motif peptides and correlate these results with cellular protein association via co-purification assays.

## Results

### G3BP1 phenylalanine residues contribute differentially to USP10 binding

The highly conserved G3BP1 NTF2 domain binds peptide motifs from USP10 (15, 28, 29). To further understand the contribution of specific residues in the G3BP1 binding groove to the interaction, we established a binding assay using fluorescence polarization. The G3BP1 NTF2 domain spanning residues 1-139 was recombinantly expressed and purified from *E. coli*. Using a USP10-derived octameric reporter peptide (*F\**-YI**FGDF**SP; *F\**, fluorescein-aminohexanoic acid tag) containing the core FGDF motif, a binding isotherm was determined for WT-NTF2 (Figure 1A). The experimental isotherm was fit by non-linear least-squares to a single-site binding curve, yielding an equilibrium dissociation constant (*K*_D_) of 3.8 μM. This value is two-fold stronger than previously reported NTF2 *K*_D_ value of 7 μM for an FGDF-containing nsP3 peptide (32), and 30-fold stronger than the 115 μM value reported for a DSGFSFGSK peptide (27). Having established a robust and repeatable assay using several different batches of NTF2, we next sought to test the effects of mutation on binding affinity. Residues F15, F33, and F124 in NTF2 are reported to play a role in coordinating the FGDF peptide from USP10 (17, 26, 32), so we focused our investigation on these residues, mutating resides F15, F33, and F124 to tryptophan (29, 32), singly and in combinations. Additional constructs were created to test other amino-acid mutations for these three residues, including alanine and tyrosine, but most of these constructs had low expression or produced insoluble protein when introduced in combination, and thus were not included in our analyses. Substitutions of tryptophan for F15 or F33 yielded 3.5- and 7.5-fold decreases in affinity, respectively (Table 1). By itself, the F124W mutant caused no significant change in binding affinity. This agrees with multiple reports that G3BP1 harboring a F33W mutation is deficient in USP10 complex formation, whereas F124W has no effect on complex formation (26, 31, 32)

**Figure 1.**
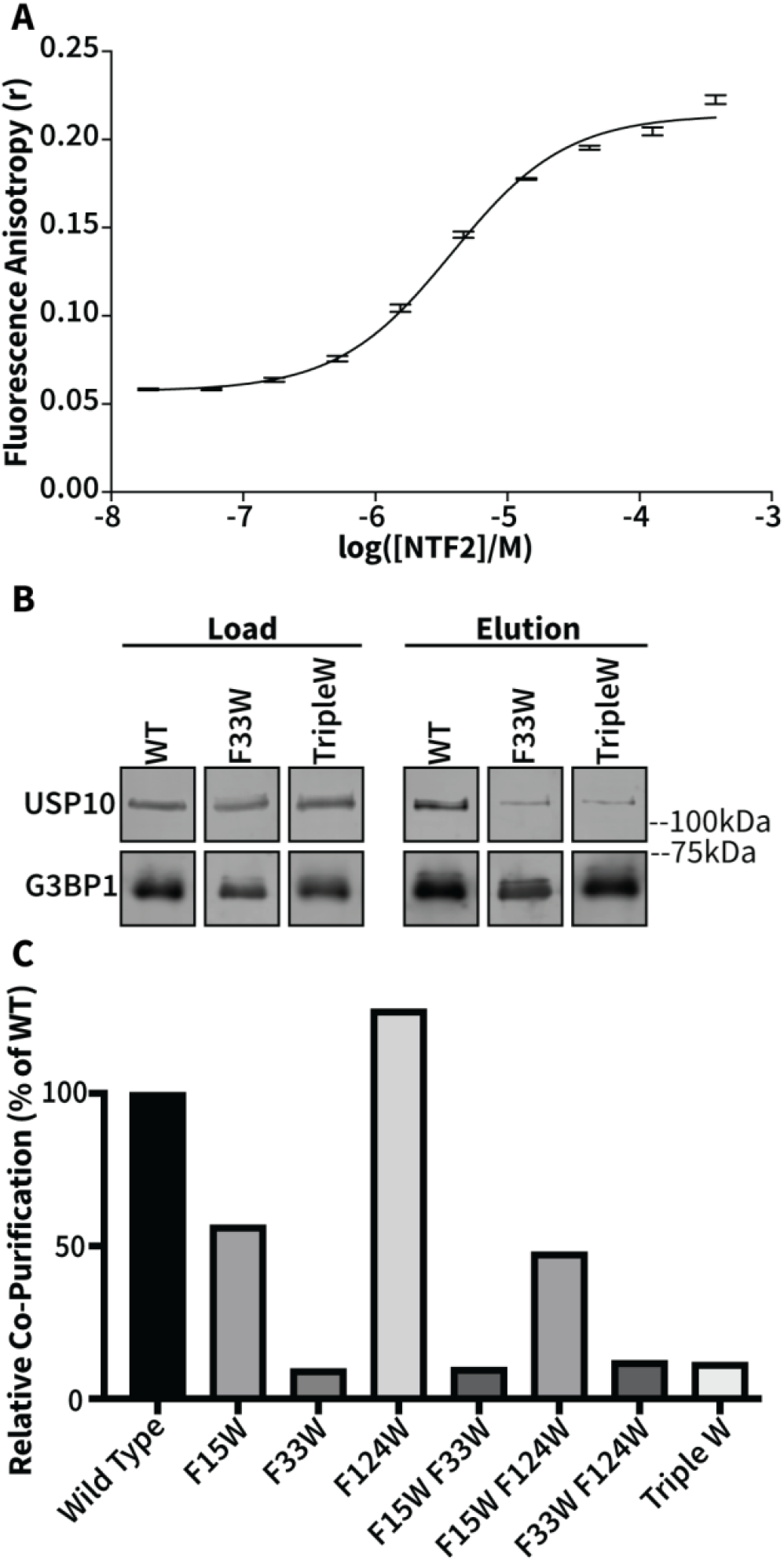
Individual equilibrium dissociation constants and co-purification of G3BP1:USP10 interactions. (A) An NTF2:USP10 binding isotherm was determined from fluorescence polarization assays. Fluorescence anisotropy data were collected using increasing concentrations of G3BP1 NTF2 in the presence of 30 nM reporter peptide derived from the USP10 FGDF binding motif (Fluorescein-aminohexanoic acid[*F\**]-YIFGDFSP). Data are shown as mean ± standard deviation of n=3 experiments. Average values were fit using a non-linear least-squares algorithm to determine *K*_D_. (B) Expi293 cells were transfected with plasmids expressing His-G3BP1 WT or mutant constructs, and lysates were subjected to affinity purification to recover His-tagged G3BP1. Equal fractions of load (left) and eluates (right) were run on SDS-PAGE followed by immunoblotting with anti-G3BP1 (bottom) and anti-USP10 (top) antibodies. (C) Immunoblots were analyzed via Image Studio, and mutants were compared to WT. Data represent the mean of n=2 experiments.

**Table 1.**
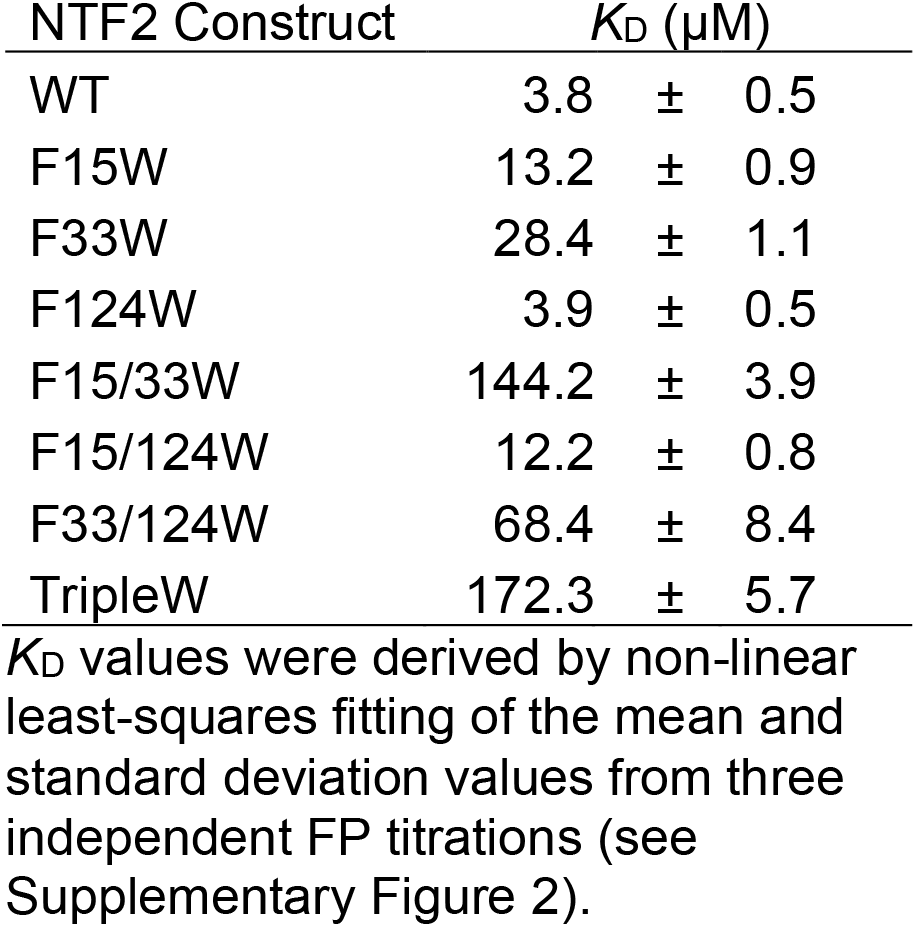
USP10:G3BP1-NTF2 Affinities.

Next we created double and triple mutants to see if the effects of the single substitutions were additive. NTF2 F15/33W and F33/124W exhibited significant increases in *K*_D_ values compared to either of the component single mutants. However, NTF2 F15/124W was only modestly worse than F15W alone. The most potent mutant was NTF2 TripleW with a *K*_D_ of 172.3 μM, although it was only modestly worse than F15/33W. These data suggest that the contribution of F124 to USP10 peptide binding is much less than might be expected based on earlier co-crystal structures (17, 26). Comparison of the F33/124W double mutant to the F33W single mutant reveals a potential context-dependent contribution of F124W in the presence of F33W. However, comparison of TripleW vs. F15/33W shows that this role of F124W is largely lost in the presence of F15W.

Our FP experiments revealed that we could manipulate the equilibrium dissociation constant of NTF2 and USP10 peptide across almost two orders of magnitude using specific tryptophan substitutions. However, we also wanted to test the effect these mutations would have on the ability of full-length versions of G3BP1 and USP10 to interact and form complexes in a cellular context. We overexpressed His-tagged full-length G3BP1 in Expi293 cells, performed immobilized metal-affinity purification, and noticed that the same molecular weight bands were co-purifying across many optimization trials (Supplementary Figure 1). We performed Western Blot analysis on the first fraction and determined that USP10 and Caprin1 were co-purifying with G3BP1 (Figure 1B). We decided to use this overexpression and affinity capture system to assay for binding stability of G3BP1 mutants. Substitution of tryptophan for F33 caused a ∼90% reduction in USP10 co-purification as compared to WT (Figure 1C). Any double and triple mutant harboring a F33W mutation (F15/33W, F33/124W, and TripleW) had a similar reduction in USP10 binding ability as compared to F33W alone. F15W alone caused a ∼44% reduction in USP10 co-purification, and when combined with F124 substitution in the F15/124W the reduction was increased to ∼53%. The F124W single mutant displayed a modest increase in USP10 co-purification as compared to WT. Since F124W does not significantly disrupt the equilibrium dissociation constant for USP10 as seen in the FP assay, overexpression of full length F124W should retain the full ability to bind and interact with USP10. When the results from these separate experiments are compared, it is evident that changes in binding affinity of G3BP1 NTF2 domains for the USP10 FGDF peptide correlate with changes in full length G3BP1:USP10 complex stability (Figure 2). Residues F15, F33, and F124 each help to stabilize the interaction with USP10, but with significantly different contributions.

**Figure 2.**
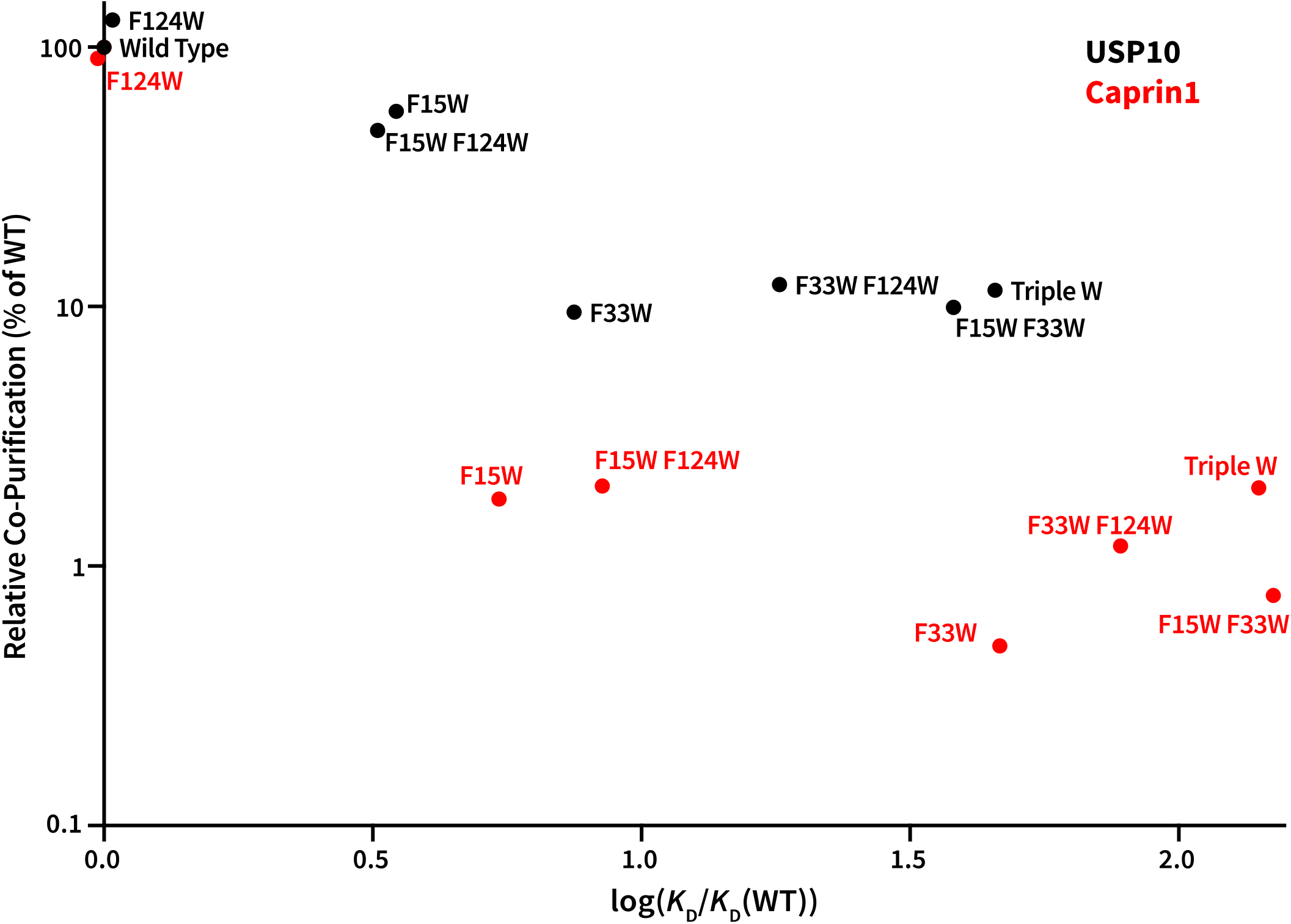
Comparison of relative co-purification of endogenous USP10 or Caprin1 versus corresponding equilibrium dissociation constants for G3BP1 mutants. Full-length His-G3BP1constructs were overexpressed and affinity purified, and relative co-purification percentages were calculated as described in Figures 1 and 3. Equilibrium dissociation constants (*K*_D_) were calculated by fluorescence polarization assays using G3BP1 NTF2 (1-139) and USP10 or Caprin1 reporter peptides. Each point represents the relative co-purification percentage and *K*_D_ for a G3BP1_mutant_:USP10 (black) or G3BP1_mutant_:Caprin1 (red) interaction. See also Figures 1 and 3.

### Caprin1 is more sensitive than USP10 to G3BP1 NTF2 modifications

Caprin1 competes with USP10 in binding the NTF2 domain, however Caprin1 lacks an FGDF motif (29). Solomon *et al*. reported that Caprin1 contains a conserved binding motif, FIQDSMLD, spanning residues 372-379 (28). Given the differences in peptide recognition sequences, we wanted to determine whether the three phenylalanine residues (F15, F33, and F124) identified as USP10 modulators also participate in binding and coordinating the Caprin1 peptide. We created a Caprin1-derived dodecamer reporter peptide (*F\**-YN**FIQDSMLD**FE) to be used in FP assays. Using recombinant NTF2 and the Caprin1 reporter peptide, we generated a binding isotherm that could be fit to a single-site binding curve, yielding an estimate of the equilibrium dissociation constant of 1.7 μM (Figure 3A). This is approximately 2.2-fold stronger than the *K*_D_ calculated for USP10, which disagrees with a recent publication from Schulte *et al*. that reported a stronger interaction between USP10:NTF2 than between Caprin1:NTF2 in ITC experiments (30). One possible explanation for this discrepancy is the pH: Schulte *et al* used pH 7.5, whereas we used pH 8.5 in our experiments. G3BP1 residues H31 and H62 have been implicated in Caprin1 binding and have predicted p*K*_a_-values of 6.1 and 6.2, respectively. Depending on the local pKa of the histidine side chains, they could be differentially protonated at pH 7.5 or pH 8.5. Alternatively, this discrepancy could be explained by the construct design, as the ITC experiments used longer (∼30 aa) USP10 and Caprin1 peptides. If correct, this suggests that residues outside of canonical binding motifs may also contribute to these protein interactions. However, the affinities of the longer peptides are not substantially different from those observed with our much shorter motif peptides, suggesting that the extent of modulation by flanking residues is relatively modest.

**Figure 3.**
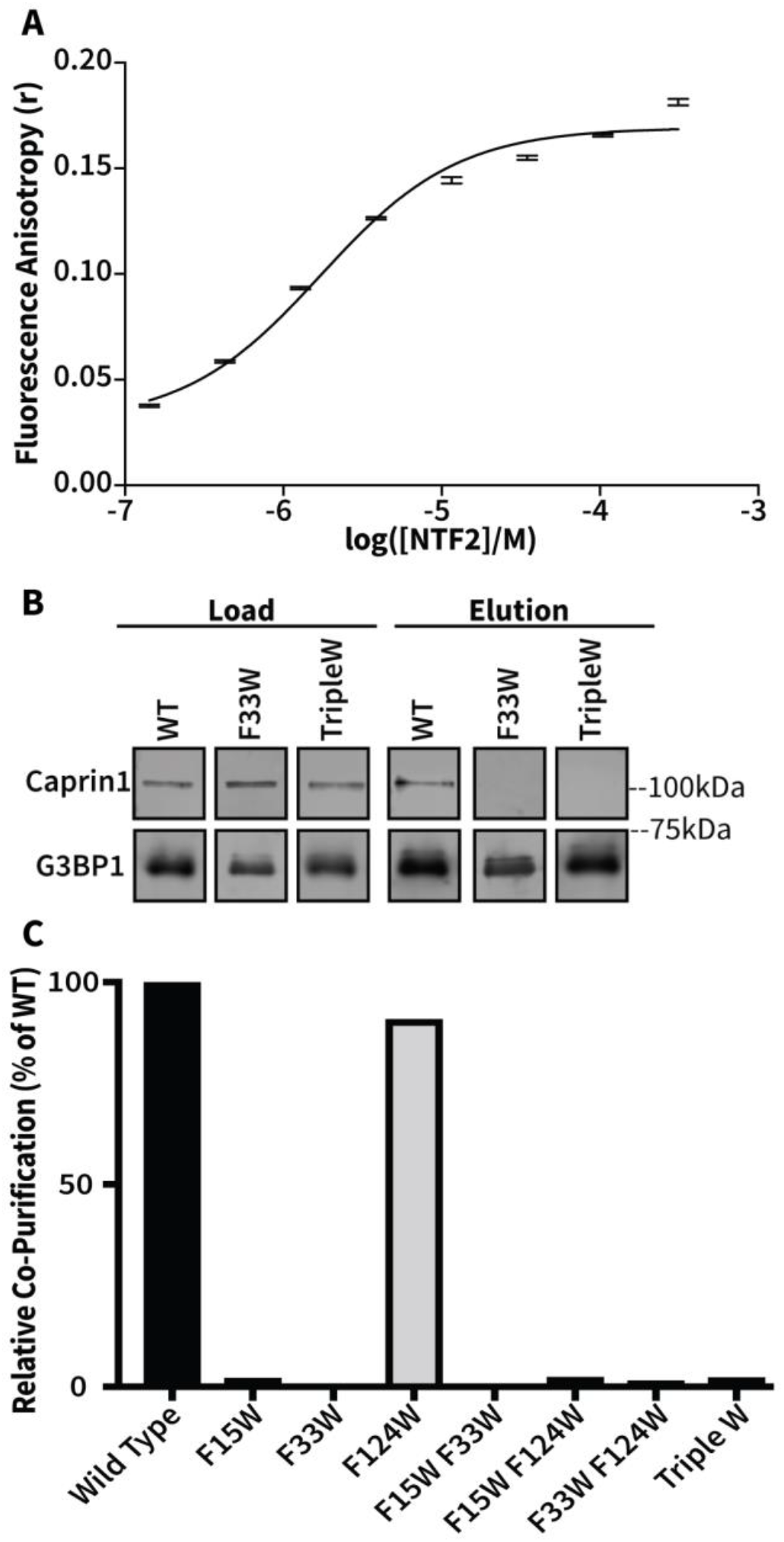
Individual equilibrium dissociation constants and co-purification of G3BP1:Caprin1 interactions. (A) An NTF2:Caprin1 binding isotherm was determined from fluorescence polarization assays. Fluorescence anisotropy data were collected, reported, and fit as described in Figure 1, using increasing concentrations of G3BP1 NTF2 in the presence of 30 nM reporter peptide derived from the Caprin1 binding motif (*F\**-YNFIQDSMLDFE). Data represent the mean ± standard deviation of n=3 experiments. (B) Expi293 cells were transfected with plasmids expressing His-G3BP1, WT or mutant, and lysates were subjected to affinity purification. Equal fractions of load (left) and eluates (right) were run on SDS-PAGE followed by immunoblotting with anti-G3BP1(bottom) and anti-Caprin1 (top) antibodies. (C) Immunoblots were analyzed via Image Studio and mutants were compared to WT. Data represent the mean of n=2 experiments.

Mutation of F15 or F33 to tryptophan caused 5.5- or 46-fold decreases in affinity, respectively (Table 2). The double mutants followed a similar trend as in the USP10 assays with F15/33W and F33/124W having a more pronounced loss of affinity as compared to the individual single mutants. F15/124W only modestly reduced the affinity as compared to F15W, further suggesting that F124 does not contribute significantly to Caprin1 peptide binding, at least in the presence of F33. Unexpectedly, the TripleW mutant exhibited slightly increased affinity as compared to F15/33W. This discrepancy may be due to relatively poor solubility of the TripleW in the FP assay. All mutations caused greater relative (fold) decreases in affinity for Caprin1 as compared to USP10.

**Table 2.**
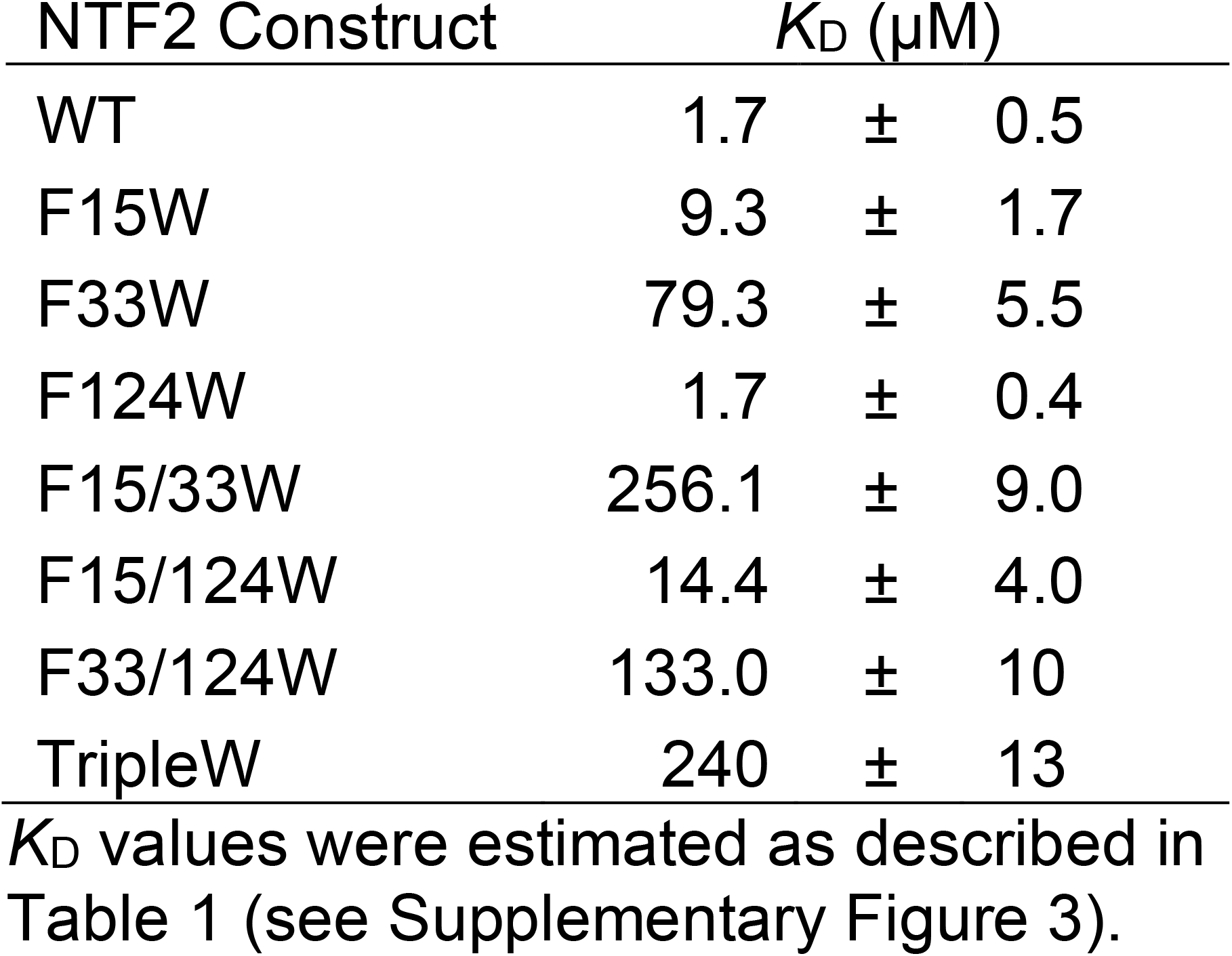
Caprin1: G3BP1-NTF2 Affinities.

Our FP assay results revealed that Caprin1 peptide binding was sensitive to mutation of residues F15 and F33. To determine the biological consequences of these mutations, we again overexpressed full length His-tagged G3BP1 mutants in Expi293 cells and assayed for binding efficiency using the affinity capture co-purification (Figure 3B). With the exception of F124W, all mutants displayed a substantial deficiency in Caprin1 co-purification, with reductions of ∼98% as compared to WT (Figure 3C). Thus, in contrast to USP10, small fold changes in *K*_D_ values caused dramatic changes in full length G3BP1:Caprin1 complex stability, such that NTF2 domains with the same experimental 1:1 affinity for Caprin1 and USP10 systematically co-purified less Caprin 1 (Figure 4).

**Figure 4.**
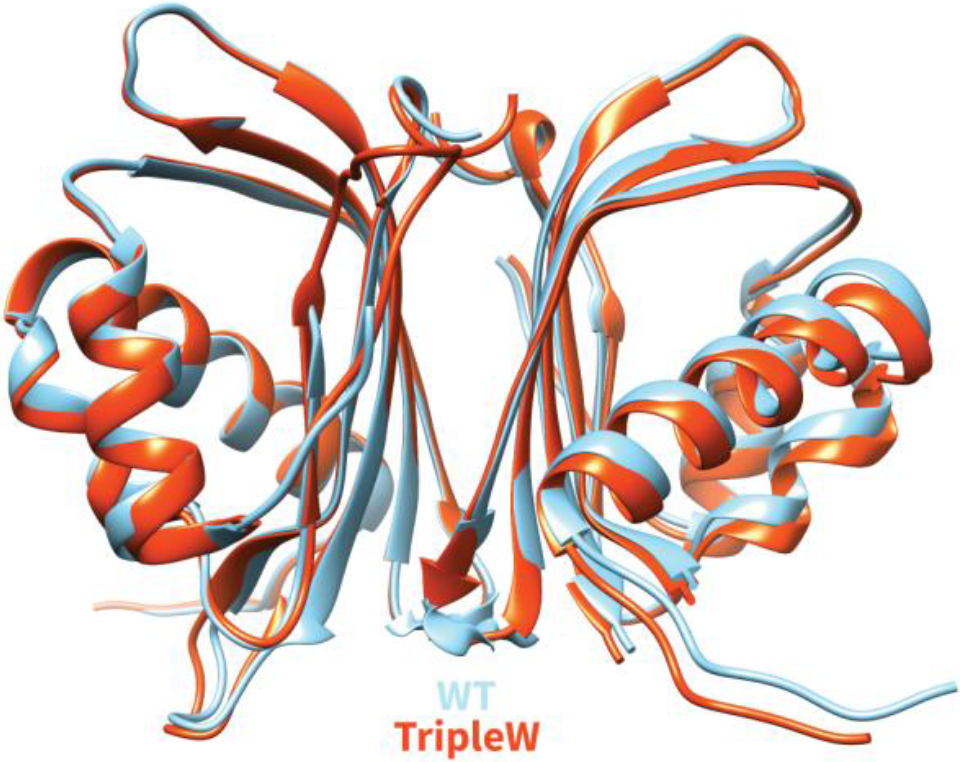
Structural conservation of the TripleW mutation. A cartoon representation of human G3BP1 NTF2-like domain TripleW (rust) is shown superimposed on G3BP1 NTF2-like domain WT (sky blue) (PDB ID 4FCJ), with all-atom calculated rmsd of 0.57 Å.

Sanders *et al*. recently found a missense mutation in NTF2 (S38F), which blocks stress granule (SG) formation and is unable to form high-affinity complexes with Caprin1 (7). Unfortunately, NTF2 S38F expressed into insoluble inclusion bodies in *E. coli*. Mutations to Ala, Gly, or Thr were well tolerated, and we tested these mutants in USP10 and Caprin1 FP assays. As expected, all three mutants decreased the affinity for Caprin1, and at similar levels to F15W (Table 3). However, in the USP10 FP assay, NTF2 S38A or S38G increased affinity for USP10 by 2.9- or 2.4-fold, respectively, whereas S38T decreased affinity by 2.4-fold (Table 3). S38 is located on a loop away from the NTF2 binding groove, so these results suggest additional G3BP1 residues contribute to peptide binding. Additionally, the data further confirm prevailing hypotheses that small changes in affinities can have dramatic changes on protein-protein interactions.

**Table 3.** Affinity of NTF2 S38 Mutants for USP10 and Caprin1.

| Mutant | $K_D$ (USP10) (μM) | | | $K_D$ (Caprin1) (μM) | | |
| --- | --- | --- | --- | --- | --- | --- |
| S38A | 1.3 | ± | 0.3 | 7.3 | ± | 1.5 |
| S38G | 1.6 | ± | 0.3 | 10.5 | ± | 2.0 |
| S38T | 9.0 | ± | 0.4 | 11.8 | ± | 0.4 |

$K_D$ values were estimated as described in Table 1 (see Supplementary Figures 2 & 3).

### TripleW can cause significant conformational changes

Several crystal structures have been published of G3BP1 NTF2 in its apo form and complexed with peptides; however, no mutant crystal structures have been published. Given the dramatic reduction in complex formation as seen with the F33W and TripleW G3BP1 mutants, we wanted to determine the stereochemical cause of the defect. A crystal structure of G3BP1 NTF2 (1-139) TripleW was determined to a resolution of 2.36 Å. The structure was solved by molecular replacement using a deposited structure of G3BP1 NTF2 (PDB ID 4FCJ). The crystal structure is in space group *P* 2_1_2_1_2_1_, with two molecules per asymmetric unit. The structure was refined to final R_work_ and R_free_ values of 22.2% and 27.1%, respectively, with excellent model geometry (Table 4).

**Table 4.** Crystal Structure Statistics.

| **Data collection & reduction** |  |
| --- | --- |
| Beamline | NSLS-II 17-ID-1 |
| Wavelength (Å) | 0.92011 |
| Space Group | $P 2_1 2_1 2_1$ |
| Unit cell parameters: |  |
| $a, b, c$ (Å) | 42.65, 71.67, 87.73 |
| $\alpha, \beta, \gamma$ (°) | 90, 90, 90 |
| Resolution<sup>a</sup> (Å) | 19.91-2.36 (2.5-2.36) |
| $R_{\text{meas}}^b$ (%) | 36.6 (231.4) |
| $\text{CC}_{1/2}^c$ (%) | 99.1 (34.9) |
| $I/\sigma_I$ | 7.02 (1.03) |
| Completeness | 99.7 (99.9) |
| Redundancy | 6.4 (6.0) |
| **Refinement** |  |
| Total number of reflections | 11543 (1793) |
| Reflections in the test set | 580 |
| $R_{\text{work}}^d/R_{\text{free}}^e$ (%) | 22.2/27.1 |
| Number of atoms: |  |
| Protein | 2204 |
| Water | 17 |
| Ramachandran plot<sup>f</sup> (%) | 95.75/4.25/0 |
| RMSD bond length (Å) | 0.01 |
| RMSD bond angle (°) | 1.07 |
| **PDB ID** | **7S17** |

<sup>a</sup>Values in parentheses correspond to the highest resolution shell.
<sup>b</sup> $R_{\text{meas}}$ : the redundancy independent R-factor, described in Diederichs and Karplus (1997); Nat. Struct. Biol. 4, 269–275.
<sup>c</sup> $\text{CC}_{1/2}$ : the percentage of correlation between intensities from random half-datasets, described in detail in Karplus and Diederichs (2012); Science 336, 1030–1033.
<sup>d</sup> $R_{\text{work}} = \sum_h |F_{\text{obs}}(h) - F_{\text{calc}}(h)| / \sum_h F_{\text{obs}}(h)$ , $h \in \{\text{working set}\}$ .
<sup>e</sup> $R_{\text{free}} = \sum_h |F_{\text{obs}}(h) - F_{\text{calc}}(h)| / \sum_h F_{\text{obs}}(h)$ , $h \in \{\text{test set}\}$ .
<sup>f</sup>Favored/allowed/outliers.

The structure of G3BP1 NTF2 TripleW could be modeled from residues 1-138 except for loop residues 48-50 and 117-123 in Chain B. Density for these loops is absent in other published apo crystal structures of G3BP1 NTF2 from the same space group. NTF2 TripleW dimerizes similarly to WT and has no major changes in backbone conformations, with an all-atom calculated root mean square deviation (rmsd) for the dimer of 0.57 Å (Figure 4).

The TripleW structure reveals major side-chain conformational changes in the binding groove. Residue 33 is dramatically twisted and shifted out of the groove when mutated to a tryptophan in Chain A (Figures 5A and 5C). In Chain B, W33 is similarly shifted out but with a smaller twist (Figures 5B and 5D). While the conformational change in W33 differs in Chain A and Chain B, both conformations are distinct from the WT F33 conformation (Supplementary Figure 4). Mutation of F33 to tryptophan generates more free energy and allows for create flexibility in the W33 side chain. The TripleW crystal structure demonstrates that W33 can adopt at least two low-energy conformations that are distinct from the WT F33 conformation. In Chain A, the large conformational change in F33W causes it to occupy the steric volume that would normally be occupied by F13 from the USP10 FGD<u>F</u> motif peptide (Figure 6A). F33W creates a similar residue-level clash with I373 from the Caprin1 FIQDSMLD target sequence (Figure 6D). In Chain B, the conformational change in W33 does not create a steric clash with either the USP10 or Caprin1 peptide but W33 occupies a larger volume than the native phenylalanine (Supplementary Figure 5). Whichever of these conformation(s) may be found in solution, the binding data confirm that accommodation of the W33 side chain imposes a free-energy cost on the binding of the peptides.

**Figure 5.**
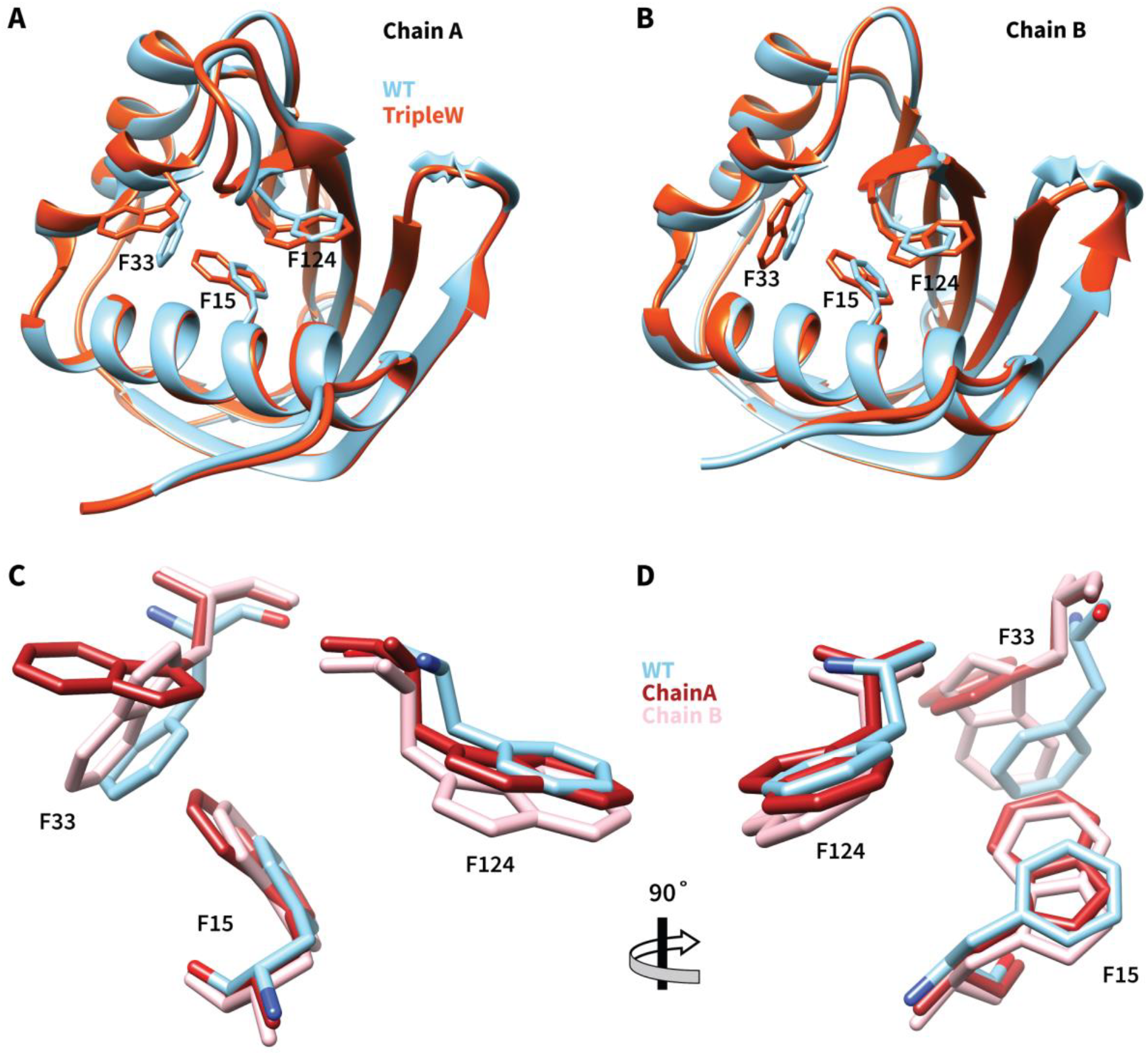
Stereochemical basis of Trp mutations in the NTF2 binding cleft. Cartoon representations of G3BP1 NTF2 Chain A (A) and Chain B (B) are shown, with TripleW (rust) superimposed onto WT (sky blue) (PDB ID 4FCJ). (C,D) Close-up stick-figure models show the three phenylalanine residues that are mutated to tryptophan in the TripleW structure. TripleW Chain A: red, TripleW Chain B: pink, WT Chain A: sky blue.

**Figure 6.**
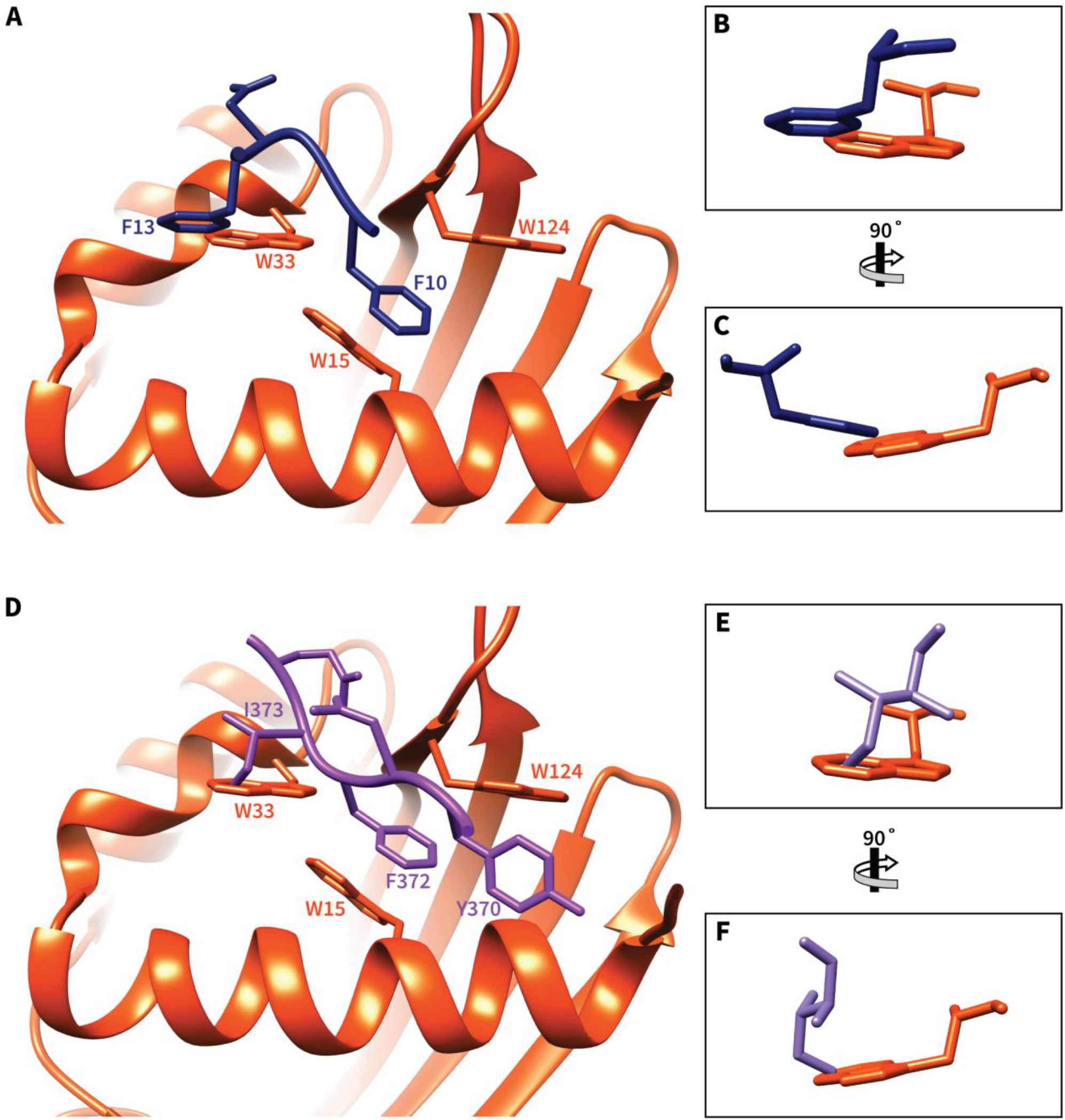
Impact of G3BP1 F33W mutation on USP10 and Caprin1 peptide binding. A cartoon representation of G3BP1 NTF2 Triple W is shown superimposed with the FGDF (A) and YNFIQD peptides (D) from the corresponding WT structures. The NTF2 TripleW structure was aligned with G3BP1 NTF2 WT co-crystal structures, and then the WT NTF2 domain was omitted from the figure. (B, C) Isolated orthogonal views of the side-chain clash between G3BP1 NTF2 W33 (rust) and USP10 F13 (navy). (E, F) Isolated orthogonal views of the side-chain clash between G3BP1 NTF2 W33 (rust) and Caprin1 I373 (purple). The following co-crystal structures were used: (A, B, C) Co-crystal structure of G3BP1 NTF2 WT with USP10 FGDF motif peptide (PDB ID 5DRV), (D, E, F) Co-crystal structure of G3BP1 NTF2 WT with Caprin1 YNFIQD motif peptide (PDB ID 6TA7).

Mutations of F15W or F124W do not significantly alter the respective side-chain conformations, such that the mutant side chains overlay with the native side chains with steric differences confined to the difference in the size of the ring systems: replacement of phenylalanine with a tryptophan does increase the side chain volume, which in turn shrinks the peptide-binding groove (Figures 5C and 5D). While F15W and F124W mutations cause changes in G3BP1 NTF2, they are much less dramatic than the changes associated with F33W. These differences likely explain why F33W has the most detrimental effect on binding affinity and complex formation.

### Mutations in USP10 FGDF motif affect G3BP1 binding

Having identified key residues in the G3BP1 NTF2 binding groove responsible for binding USP10 and Caprin1 peptides, we turned to investigating residues in USP10 and Caprin1 that influence G3BP1 binding. While a previous group published an alanine scan in a FGDF-containing nsP3 peptide (32), there has yet to be a full amino-acid substitution analysis of the proposed FGDF or FIQDSMLD core motifs. We designed a peptide array in which the residues in the core motifs are individually substituted with all 20 genetically encoded amino acids. In the USP10 microarray (Figure 7A), we chose to extend our analysis beyond the core motif to include three additional residues on the N- and C-terminal sides of the motif, so we scanned through the residues of a QYI**FGDF**SPD reference peptide. We performed the peptide array experiment at a protein concentration of 500 nM (∼9.1 μg/mL) of NTF2 protein; visualization of the amount of bound protein by antibody staining yielded variable spot intensities (Figure 7A). As expected, the core FGDF motif contributes most of the binding specificity: most substitutions in these four amino acids reduced or abrogated G3BP1 NTF2 binding. The glycine residue (G11) is the most stringent requirement of the core motif, since all substitutions, exception for alanine, dramatically reduce binding. The first phenylalanine residue (F10) tolerates tyrosine substitution, and to a much lesser extent, valine, leucine, and isoleucine. The second phenylalanine (F13) tolerates substitution to tyrosine, valine, leucine, and isoleucine better than F10, and can accommodate a tryptophan. The aspartic acid (D12) is the least strict of the core motif residues, because it can be mutated to most amino acids without dramatic changes in binding ability. The three upstream and downstream residues appear to have only modest and variable effects on G3BP1 NTF2 binding. Most mutations in these residues do not have dramatic changes in binding ability, although there may be some opportunities to enhance affinity by concerted tuning of these peripheral residues (47, 48).

**Figure 7.**
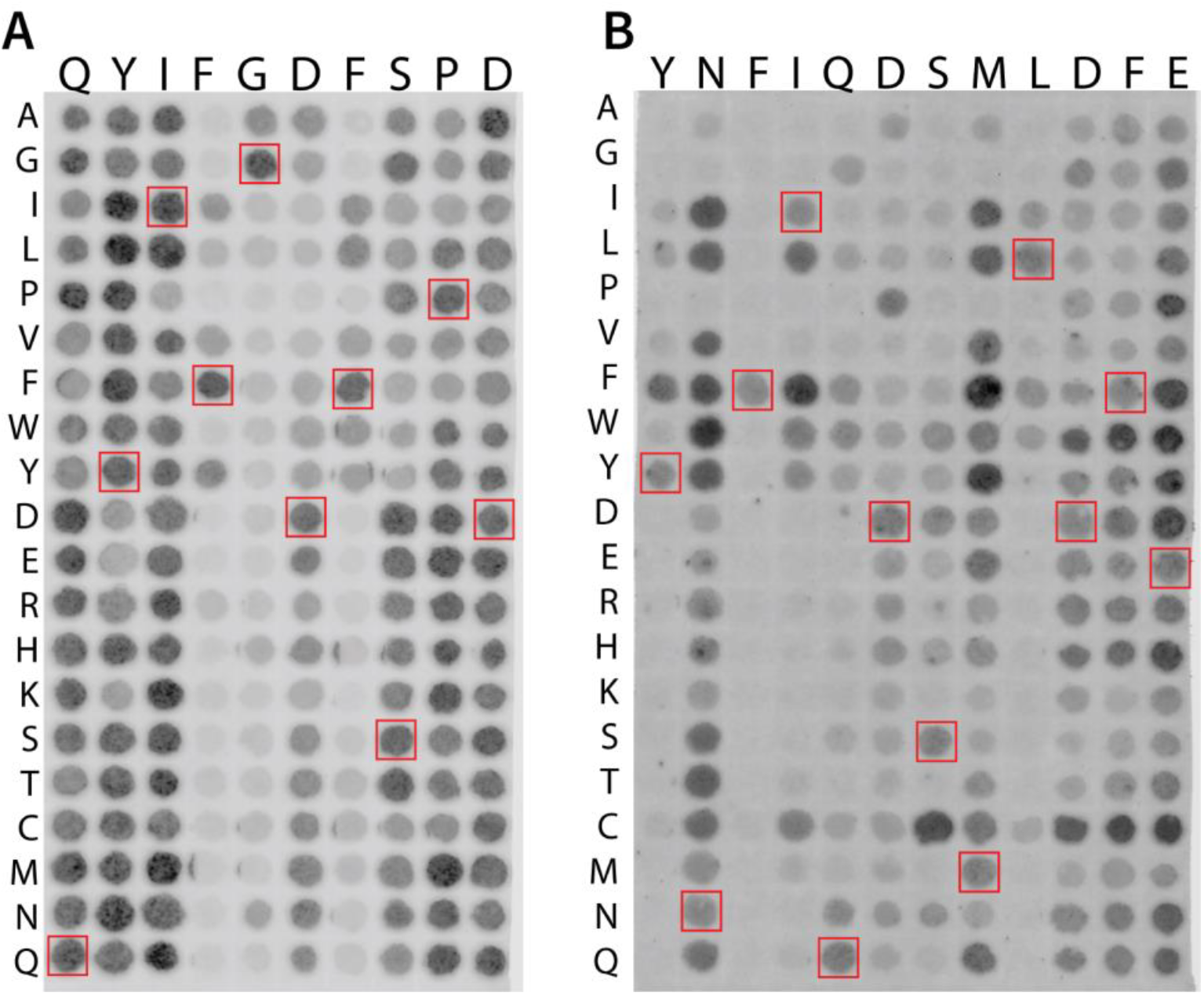
Binding of G3BP1 NTF2 to substituted USP10 and Caprin1 peptides. (A) Peptide cellulose SPOT array with single amino-acid substitutions in the USP10 binding peptide. Each spot contains the native USP10 peptide (HSP**QYIFGDFSPD**EFNQFFV) with a single amino acid in the native sequence (shown in the top row for bold residues in sequence) being replaced by the specific amino acid described by the amino-acid code in the left column. (B) Peptide cellulose SPOT array with single amino-acid substitutions in the Caprin1 binding peptide. Each spot contains the native Caprin1 peptide (QGP**YNFIQDSMLDFE**NQTLD) with a single amino acid in the native sequence (shown in the top row for bold residues in sequence) being replaced by the specific amino acid described by the amino-acid code in the left column. Red boxes highlight peptides that match the native sequence. Darkness represents bound G3PB1 NTF2 protein as detected via antibody staining protocol.

### Caprin1 has multiple residue interactions with G3BP1

In the Caprin1 peptide array, we extended our amino-acid scan to include two additional residues on the N- and C-terminal side of the previously reported core motif yielding YN**FIQDSMLD**FE as the reference sequence (28). Since our Caprin1 reporter peptide has a higher baseline affinity for G3BP1 NTF2 as calculated by FP, we incubated this array with 300 nM (∼5.5 μg/mL) of recombinant G3BP1 NTF2 (Figure 7B). The phenylalanine residue at the N-terminus of the core motif (F372) appears to be the most stringent residue in the core motif, since all substitutions completely abrogated binding to G3BP1 NTF2 (Figure 7B). Substitutions at other positions were also disruptive, as expected. Surprisingly, we also found substitutions in the core motif that resulted in higher affinity peptides. For example, both I373F and S376C exhibited dramatic increases in the amount of bound G3BP1 NTF2. Additionally, replacements of M377 with phenylalanine or tyrosine each increased binding. The conserved core motif reported by Solomon *et al*. had flexibility in three residues: F(M/**I**/L)Q(**D**/E)Sz(I/**L**)D (28). I373 substitution by leucine modestly increased affinity, where a methionine at this position ablated binding. Substitutions at the other two residues D375E and L378I modestly reduced binding. Surprisingly, Y370, which lies outside of the earlier core motif, appears to play an important role in binding: at this position all amino acids except for phenylalanine dramatically reduce binding. This suggests that the Caprin1 motif should be extended to include the residue Y370. Consistent with this proposal, mutation of N371 to alanine, glycine, or proline completely ablated binding, suggesting that it also lies within the binding motif. However, all other substitutions were well tolerated, and some substitutions (Ile, Leu, Phe, and Trp) created higher affinity peptides. Substitutions of C-terminal flanking residues F380 and E381 were well tolerated and often produced higher affinity spots. Consistent with these results, and in contrast to USP10, trimming of these motif flanking residues also created lower affinity peptides (Supplementary Figure 6). These data suggest that G3BP1 NTF2 interacts with more residues when binding Caprin1 than when binding USP10, which may account for the differential affinity we observed.

Additionally, we analyzed another reported G3BP1 interaction partner, USP13, which like USP10 has also been implicated in the post-maturational stability of CFTR (49, 50). The USP13 binding motif has not been previously reported, so we performed a scan of the N-terminal disordered region of the protein using overlapping 22mer peptides. Multiple spots contained peptides that appeared to interact with G3BP1 NTF2 (Supplementary Figure 7), including the N-terminus, as well as a cluster involving peptides spanning amino acids 73-118. These data suggest that USP13 might have a relatively long motif peptide similar to Caprin1, or alternatively, that multiple motifs might be present within USP13. Further investigations will be needed to definitively identify the USP13 binding motif(s).

## Discussion

In this work, we performed a thorough mutational analysis of the generative interactions that form G3BP1:USP10 and G3BP1:Caprin1 complexes. Using the smallest necessary components for these protein-protein interactions, the G3BP1 NTF2 domain and a motif peptide, we determined the binding affinities for multiple mutants ranging across a spectrum of two orders of magnitude. These results show that the G3BP1 NTF2-like domain has a modestly stronger affinity for Caprin1 peptide versus USP10 peptide. Conversely, we showed that binding of the longer Caprin1 peptide is more sensitive than USP10 to mutations in the G3BP1 binding groove. Mutation of G3BP1 residues F15, F33, and/or F124 caused higher fold decreases in binding affinity for Caprin1 versus USP10. Furthermore, larger amounts of USP10 co-purify with overexpressed G3BP1 NTF2 mutants with a given affinity for USP10 compared to the amounts of Caprin1 that co-purify with G3BP1 NTF2 mutants that have similar 1:1 affinity for Caprin1 (Figure 4). In agreement with previous literature, substitution of F33 for a tryptophan in the G3BP1 NTF2-like domain substantially reduces binding for both USP10 and Caprin1. However, G3BP1 F33W was able to co-purify a small amount of USP10 (∼10% compared to WT) but not Caprin1, further supporting the hypothesis that Caprin1 is more sensitive to disruption by mutagenesis than USP10, despite its higher baseline affinity. In any case, relatively modest differences in protein:peptide interaction affinities have been associated with significant differences in the assembly of protein complexes. For example, the Ki-67 and RepoMan phosphatase regulatory proteins exhibit strong selectivity for protein phosphatase 1 (PP1) γ isoform over the α isoform, based on a roughly four- to six-fold difference in *K*_D_ arising primarily from a single amino-acid difference (51).

Co-purification has classically been used in non-affinity chromatography schemes to purify tagless proteins from cells (52, 53). Proteins co-purified using these schemes do not necessarily interact or form complexes directly, so additional experiments were necessary to prove protein-protein interactions. In our case, the direct interaction has already been established biochemically and crystallographically using purified components. Recent reports, however, have shown the benefits of affinity chromatography in co-purifying protein complexes such as antibody stabilized proteins, RNA-protein complexes, and cytochrome supercomplex (54–56). The homogeneity of these purified complexes has been confirmed by biophysical methods such as small-angle X-ray scattering (SAXS) and protein crystallography (8, 54). G3BP1, USP10, and Caprin1 form protein complexes that are present within SGs that participate in liquid-liquid phase separation (LLPS) (7–9). Several reports have detailed the fluidity and reversibility of SGs (7–9); however, the G3BP1 complexes persisted through a strenuous purification scheme. This suggests that while the larger SG bodies may be fluid, the protein complexes within them may be unexpectedly persistent. Purification of protein complexes via metal affinity allows us to study them *in vitro* without the need to reconstitute the complex using recombinant versions of each individual component. And while not reported here, these purified protein complexes have been useful in development of additional assays. Given the complexity of our system, it was beneficial to use affinity co-purification rather than the more common co-immunoprecipitation. While we report our co-purification data as in terms of percentage of WT recovery, we are effectively measuring the off-rate of these protein-protein interactions, which appears to be very slow, consistent with multivalency. The data reported here provide additional evidence that affinity co-purification is a useful tool for *in vitro* investigations of large proteins complexes.

Based on prior data, we were somewhat surprised by the results of the FP assay and co-purification experiments using the F15W mutant. In the FP assays, G3BP1 NTF2 F15W had calculated *K*_D_ values of 11.6 μM and 9.3 μM for USP10 and Caprin1, respectively. These values are similar and are each less than one order of magnitude different from WT, so we expected reduced, but not abolished, USP10 and Caprin1 co-purification. Indeed, the longer Caprin1 binding motif (FIQDSMLD) might have been expected to be more stable to disruption by a single amino-acid substitution. However, the opposite result was observed. While USP10 co-purification was reduced by ∼50% compared to WT, the G3BP1 F15W mutation almost entirely abolished Caprin1 co-purification.

One possible explanation is that the off-rate of the larger Caprin1 binding peptide may be faster, even if the affinity is similar. Previous crystal structures have demonstrated the importance of hydrophobic interactions and π-stacking in the G3BP1 groove for USP10 and Caprin1 peptide binding (17, 26, 30). As a result, this disjunction between USP10 and Caprin1 binding may well reflect differences in the motif peptides, since the USP10 FGDF motif contains two phenylalanines, whereas the Caprin1 FIQDSMLD motif contains only one. G3BP1 F15 participates in parallel-displaced π-stacking with USP10 F10 and Caprin1 F372. When mutated to a tryptophan, residue 33 occupies a larger hydrophobic volume with a slightly adjusted side chain (Figure 5). This change sterically occludes the positioning the USP10 and Caprin1 peptides (Figure 6), most likely accounting for the observed reduction in affinity. The impact on the USP10 off-rate may be less dramatic, since it has a second phenylalanine, F13, that can still form π-stacking interactions with G3BP1 F33. During binding, USP10 F10 and Caprin1 I373 occupy the same position in the G3BP1 binding groove. Interestingly, in the Caprin1 microarray experiment, I373F created a higher affinity peptide. Thus is it possible that Caprin1 I373F could rescue Caprin1 co-purification in the G3BP1 F15W overexpression system.

The literature describing G3BP1:USP10 and G3BP1:Caprin1 complexes has primarily focused on mutational and functional analyses of G3BP1 residues and binding interfaces (7–9, 29, 30). We also wanted to understand the contributions from USP10 and Caprin1 in complex formation. Using a systematic amino-acid scan, we have confirmed that the USP10 FGDF motif is the smallest necessary component for G3BP1 NTF2 binding (32). As expected, most substitutions in the core FGDF motif created lower affinity peptides, while substitutions in the upstream and downstream residues modulated affinity, but were mostly well tolerated (Figure 7A). In particular, substitution of flanking residues Y8 and I9 has the potential to create higher affinity peptides. In the Caprin1 array, we saw that most substitutions in the FIQDSMLD motif created lower affinity peptides (Figure 7B), again confirming the importance of the previously reported motif (28). However, substitutions I373F, S376C, M377F, and M377Y created higher affinity peptides. Given that I373 sits similarly to USP10 F13, it is unsurprisingly that replacement with a phenylalanine creates a higher affinity peptide. Similar to the USP10 peptide array, upstream and downstream amino-acid substitutions often created higher affinity peptides; however, flanking residues have a greater role in Caprin1 binding (Supplementary Figure 6). In particular, Y370 appears to play a critical role in G3BP1 binding since it only tolerated substitution to phenylalanine (Figure 7B), and binding was abolished when it was removed (Supplementary Figure 6).

G3BP1:USP10 and G3BP1:Caprin1 complex formation are associated with several disease (57–60). Our peptide array studies suggest that multiple substitutions in the USP10 and Caprin1 binding motifs can create higher affinity peptides. Combining several of these substitutions in one peptide should have additive effects to create significantly higher affinity competitor peptides. Interestingly, the interactions of G3BP1, USP10, and Caprin1 can be cooperative or competitive within the context of multivalent SGs (7–9). Mutations in the G3BP1 NTF2 binding site had disparate effects on Caprin1 and USP10, and their motifs are significantly different in length and composition. As a result, peptide inhibitors based on these different motifs may have differential effects on the composition and stability of the resulting complexes in cells, which can be readily assessed using the co-purification assay described here.

Depending on which patterns are observed, it may ultimately prove therapeutically useful to target one of both component interactions selectively. Given that the necessary component for complex formation is NTF2 and short motif peptides, we could easily adapt our FP assay to perform high-throughput screens for novel inhibitors. Indeed, we have previously shown in a separate protein-peptide interaction that our medium-throughput FP assay is easily translated to high-throughput screening studies (61). Currently, there are two published G3BP1 targeted peptide therapies including GAP161 (22), which blocks RasGAP association, and GAP159 (62), which inhibits G3BP1 expression. Resveratrol, an anticancer agent, has been shown to bind the G3BP1 NTF2-like domain and induce apoptosis via p53 activation (31, 63). While these therapies are targeted towards cancer progression and metastasis, they do support the concept of targeting G3BP1 and its complex formation as advantageous and novel drug targets.

Benefits arising from the ability to inhibit or control G3BP1 complex formation are observed in nature as well. During viral attack, infected cells will activate stress granule formation to block mRNA processing and halt viral growth (64). As an essential component of stress granules, G3BP1 plays an important role in the antiviral response. Viruses use the FGDF-containing nsP3 to bind G3BP1 and sequester it away from stress granules, thereby inhibiting stress granule formation and allowing viral growth (17). G3BP1 NTF2-like domain has been reported to interact with the nsP3 protein from many viruses including SARS-CoV-2, Old World alphavirus, Semliki Forest virus (SFV), Sindbis virus (SINV), Herpes Simplex virus, and chikungunya (24, 32, 65–69).

Finally, USP10 is a deubiquitinase that plays an important role in protein homeostasis and recycling. One target of USP10 is CFTR, a chloride transporter that helps maintain ion balance across epithelial cells in the lungs and other tissues (36). As part of its normal lifecycle, CFTR is retrieved from the plasma membrane into endosomes for peripheral quality control (70). USP10 deubiquitinates CFTR in endosomes, thereby increasing the probability of endocytic recycling, which in turn helps to maintain the abundance of CFTR at the plasma membrane. In lung infections caused by *Pseudomonas aeruginosa*, a virulence factor called Cif is secreted from the bacteria in outer membrane vesicles, and is able to trigger a reduction in plasma membrane CFTR levels (20, 71). Cif stabilizes a G3BP1:USP10 complex in lung cells which renders USP10 inactive and thus unable to deubiquitinate CFTR (20, 36, 71). Knockdown of G3BP1 blocks the Cif effect suggesting that USP10 expression and activity is protective for CFTR and ion homeostasis during *Pseudomonas aeruginosa* infections (20). Thus, while it is not known how Cif drives G3BP1:USP10 complex formation, inhibitors of the interaction could also help to neutralize Cif-facilitated CFTR degradation.

The data reported here will facilitate future experiments on the roles of G3BP1, USP10, and Caprin1. We have identified several mutations in the three proteins that alter binding affinity by large and small amounts. As we have shown, small changes in affinity can have dramatic effects on the cell biology of the system, so we can use these mutations to selectively manipulate the G3BP1-USP10-Caprin1 system. This will help deconvolute the intricate interaction network surrounding these three proteins and help address unanswered questions in disease contexts including bacterial virulence, antiviral response, innate immunity, and neurodegeneration.

## Materials and Methods

### Cloning, protein expression, and purification

The pNIC28-G3BP1-NTF2 vector was graciously provided by Dr. Gerald McInerney (Karolinska Institutet, Stockholm, Sweden). Mutants were created via side directed mutagenesis using primers designed on NEBaseChanger and Q5 site-directed mutagenesis kit following manufacturer’s directions (New England BioLabs). BL21(DE3) cells were transformed with plasmid and grown on LB (72) + 50 µg/mL Kanamycin agar plates at 37°C overnight. Transformants were used to inoculate 10 mL LB + 50 µg/mL Kanamycin broth and grown at 37°C overnight. Overnight cultures were used to inoculate 1 L LB + 50 µg/mL Kanamycin broth in non-baffled 2 L flasks and allowed to grow at 37°C with shaking. Once the OD_600_ of the cultures reached ∼0.8, the cultures were induced with 0.1 mM IPTG and transferred to 30°C with shaking for 4 hours. Cultures with mutant G3BP1 NTF2 were expressed at 16°C with shaking overnight (∼18 hours). After expression, cultures were pelleted in a JLA-9.1000 rotor at 4,500 rpm for 15 min at 4° C. Pellet was resuspended in lysis buffer (500 mM NaCl, 20 mM Tris pH 8.5, 2 mM MgCl_2_) supplemented with Pierce universal nuclease at 25 units/mL and Roche cOmplete EDTA-free Protease Inhibitor Cocktail tablets. Cell lysis was carried out using an M-110L microfluidizer (Microfluidics) in 3 passes at ∼18 kpsi. Lysate was spun down for 1 hour at 40K rpm at 4°C in a Type 45 Ti rotor. 5 mL of HisPur Ni-NTA Resin (Thermo Scientific) was washed with 25 mL wash buffer (500 mM NaCl, 20 mM Tris, pH 8.5). The clarified cell lysate was supplemented with 20 mM imidazole, pH 8.5, and then incubated with Ni-NTA resin with gentle stirring for 1 hour at 4°C. The mixture was returned to room temperature and passed through a gravity flow column. The protein-bound resin was washed four times with 25 mL of wash buffer supplemented with 80 mM imidazole, pH 8.5. The protein was eluted with 500 mM NaCl, 20 mM Tris pH 8.5, and 500 mM imidazole, pH 8.5. Eluates were immediately diluted 1:1 with 800 mM NaCl and 20 mM Tris, pH 8.5 then pooled for dialysis at room temperature into 500 mM NaCl and 20 mM Tris, pH 8.5. After at least an hour of dialysis, purified recombinant TEV was added at a ratio of 1:20 and left at room temperature overnight. The cleaved protein was passed through HisPur Ni-NTA resin equilibrated with wash buffer. The flow through was collected, concentrated, and loaded on a HiLoad 26/600 Superdex 200 pg size-exclusion chromatography column (GE Healthcare). Protein was eluted using a buffer of 150 mM NaCl and 20 mM Tris pH 8.5 at the expected molecular mass (Supplementary Figure 8).

Full-length G3BP1 was expressed and purified from Expi293F cells (Life Tech) as detailed here. The pCMV-His-G3BP1 vector was purchased from SinoBiolgical. Mutants were generated using the same site-directed mutagenesis protocol as detailed above. Expi293F cells were grown in Expi293 Expression Media at 37 °C, 8% CO_2_ with shaking at 125 rpm in a disposable plastic Erlenmeyer flask (non-baffled). On the day of transfection, cell density was between 4-5 × 10^6^ cells/mL with >95% viability as determined using a TC20 automated cell counter (Bio-Rad). Cells were diluted with warm Expi293 Expression Media to 2 × 10^6^ cells/mL in 42.5 mL. 5 µg of plasmid was added to 2.5 mL Opti-MEM Reduced-Serum Medium (Gibco). 15 mg of sterile PEI was added to 2.5 mL Opti-MEM Reduced-Serum Medium. PEI solution was added to DNA solution, inverted several times, and incubated for 20 minutes at room temperature. DNA:PEI solution added to flask of Expi293 cells then returned to incubator at 37°C, 8% CO_2_ with shaking at 125 rpm. After 16-20 hours, 25 µL of 100 mM sterile-filtered valproic acid (Sigma) and 2.5 mL of 100 mM sterile-filtered sodium propionate (Sigma) in Expi293TM Expression Medium were added to the flask. 72 hours after transfection, cells were gently pelleted, and then resuspended in lysis buffer (500 mM NaCl, 20 mM Tris pH 8.5, 1% [*v/v*] IGEPAL, 2 mM MgCl_2_) supplemented with Pierce universal nuclease at 25 units/mL and Roche cOmplete EDTA-free Protease Inhibitor Cocktail tablets. Conical tubes were placed on gentle rotator for 30 min at 4°C. Clarification and elution followed the same protocol as above. After elution, samples from first eluate were collected, boiled in SDS sample buffer, and run on a 10% SDS acrylamide gel. Proteins were transferred to a PVDF membrane at 95 V for 2 hours at 4°C. Membranes were blocked for 1 hour at room temperature in Odyssey Blocking Buffer (LiCor). Primary antibodies (rabbit polyclonal USP10 antibody, Bethyl; rabbit polyclonal Caprin1 antibody, Proteintech; mouse monoclonal G3BP1 antibody, Santa Cruz) were diluted into Odyssey Blocking Buffer with 0.2% (*v/v*) Tween 20. Membranes were incubated with primary antibody solutions overnight at 4°C. Membranes were washed with Tris-buffered saline, 0.1% (*v/v*) Tween 20 (TBST) then incubated for 1 hour in IRDye secondary antibody (LiCor) in Odyssey Blocking Buffer with 0.2% (*v/v*) Tween 20 + 0.01% (*w/v*) SDS. Membranes were washed with TBST multiple times, and then imaged with the Odyssey CLx Imager (LiCor).

### Fluorescence polarization

Fluorescence polarization assays were performed following lab established protocols (73). Briefly, experiments were performed using G3BP1 NTF2 protein and performed in triplicate. Biomatik synthesized the reporter peptides used for all FP experiments: fluorescein-aminohexanoic acid (*F\**)-YIFGDFSP (USP10) and *F\**-YNFIQDSMLDFE (Caprin1). All experiments were performed in stock buffer (150 mM NaCl, 20 mM Tris, pH 8.5) supplemented with 30 µM thesit and 0.1 mg/mL IgG. Protein was serially diluted into stock buffer containing 30 nM reporter peptide then transferred to a 384-well plate.

Protein-reporter was allowed to incubate at room temperature for 30 min. Plates were scanned on a Synergy Neo2 multi-mode plate reader (BioTek) and the anisotropy was measured. A non-linear least-squares algorithm was used to fit the merged anisotropy data and calculate *K*_D_. Anisotropy and intensity values were analyzed manually to confirm no effect due to light scattering.

### Crystallography

G3BP1 NTF2 TripleW crystals were obtained by vapor diffusion in a sitting drop composed of 200 nL of 1.6 mg/mL protein in 100 mM NaCl, 20 mM Tris, pH 8.5 and 200 nL of well solution (20% [*w/v*] PEG 8000, 100 mM HEPES pH7.5) and equilibrated by vapor diffusion with 50 µL of well solution. The drop was set up using a NT8 drop setter (Formulatrix) and the well solution was from the Wizard Classic I&II High-Throughput Screen (Molecular Dimensions). The tray was incubated and imaged using Rock Imager and Rock Maker instrumentation (Formulatrix). The crystal was harvested and immediately flash cooled in liquid nitrogen. The oscillation data collection was performed at the National Synchrotron Light Source II beamline 17-ID-2 (FMX) equipped with an Eiger 16M detector at 100 K, an oscillation range of 0.2° per frame, a total of wedge of 180°. The diffraction images were processed using XDS (74). The R_free_ set was generated from 5% of the reflections in thin resolution shells using the Phenix (75) reflection file editor. Initial phases were generated by Phaser (76) via molecular replacement using G3BP1 NTF2 WT (PDB ID 4FCJ) as a search model. Iterative automatic and manual refinement were performed using Phenix and Coot (75, 77).

### Peptide Array

The 600 peptide SPOT cellulose array was generated by the Biopolymers & Proteomics Core Facility at the Koch Institute for Integrative Cancer Research of Massachusetts Institute of Technology. An Intavis SPOT synthesis peptide arrayer system was used to make the array.

The peptide array was hydrated with methanol for 10 min, then washed three times with TBST, and then blocked for 2 hours in Odyssey Blocking Buffer. The array was incubated with His-tagged G3BP1 NTF2 in blocking buffer with 0.2% (*v/v*) Tween 20 for ∼20 hours at 4°C, and then washed three times with TBST. G3BP1 was detected using a His-tag antibody (Santa Cruz) followed by IRDye secondary antibody in Odyssey Blocking Buffer with 0.2% (*v/v*) Tween 20 + 0.01% (*w/v*) SDS. Arrays were scanned on an Odyssey CLx Imager.

### Accession Numbers

The coordinates and structure factors for the G3BP1 NTF2-like domain TripleW structure are available at the Protein Data Bank under PDB ID: 7S17.

## Funding

This work was supported by National Institutes of Health grants R01-AI091699, P20-GM113132, and P30-DK117469, training grant T32-HL134598, and beamline support P30-GM133893. Additional beamline funding was provided by US Department of Energy contracts DE-SC0012704 and KP1607011. The content is solely the responsibility of the authors and does not necessarily represent the official views of the National Institutes of Health or the US Department of Energy.

## Acknowledgments

We would like to thank Drs. Babak Andi, Martin Fuchs, and Vivian Stojanoff at FMX for support of diffraction experiments, Dr. Noor Taher for advice and suggestions regarding data processing and structure refinement, Dr. Kelli Hvorecny for mentorship and advice, Dr. Andreia Verissimo and Dr. Angela Kull for the support of the bioMT core facilities, Dr. Bradley Bartholomai and members of the Madden and Stanton lab for helpful discussions.

## Notes

### Competing Interest Statement

The authors have declared no competing interest.

