## Supplementary Figures for "Stereochemistry of transient protein-protein interactions in a signaling hub: exploring G3BP1-mediated regulation of CFTR deubiquitination"

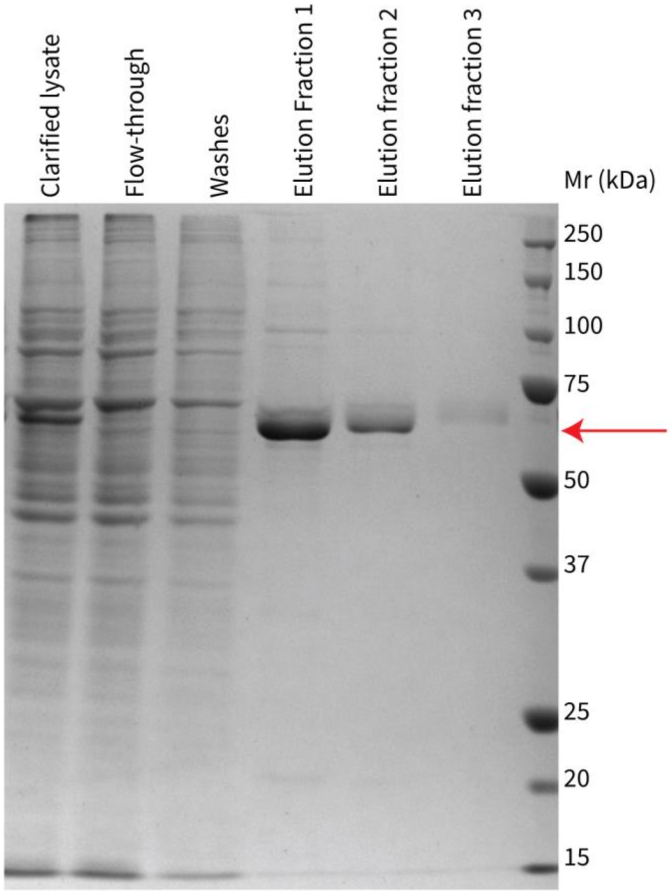


**Supplementary Figure 1. Full-length His-G3BP1 affinity purification.** A Coomassie-stained SDS-PAGE gel shows purification of full-length His-tagged G3BP1 using Ni-NTA resin. Lane identifications are shown at the top of each lane. Arrow shows expected position of G3BP1.

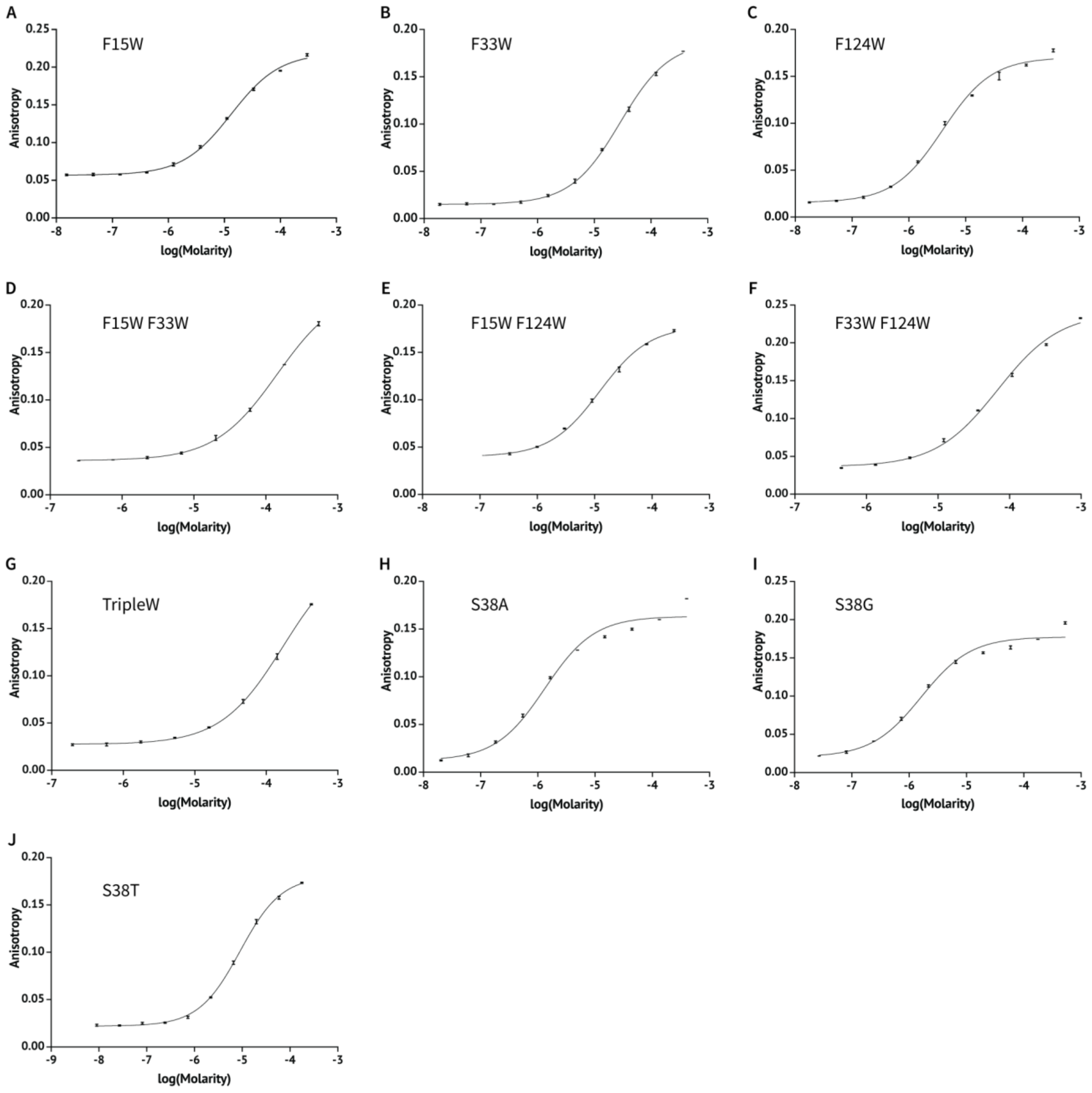


**Supplementary Figure 2. The affinities of mutant G3BP1 NTF2 domains for USP10.** Fluorescence anisotropy data were collected using increasing concentrations of G3BP1 NTF2 in the presence of 30 nM reporter peptide derived from the USP10 FGDF binding motif (*F**-YIFGDFSP). Data represent the mean and standard deviation of n=3 experiments. For each construct, the average from three independent titrations was fit using a non-linear least-squares algorithm as summarized in Table 1. (A) F15W, (B) F33W, (C) F124W, (D) F15/33W, (E) F15/124W, (F) F33/124W, (G) TripleW, (H) S38A, (I) S38G, and (J) S38T.

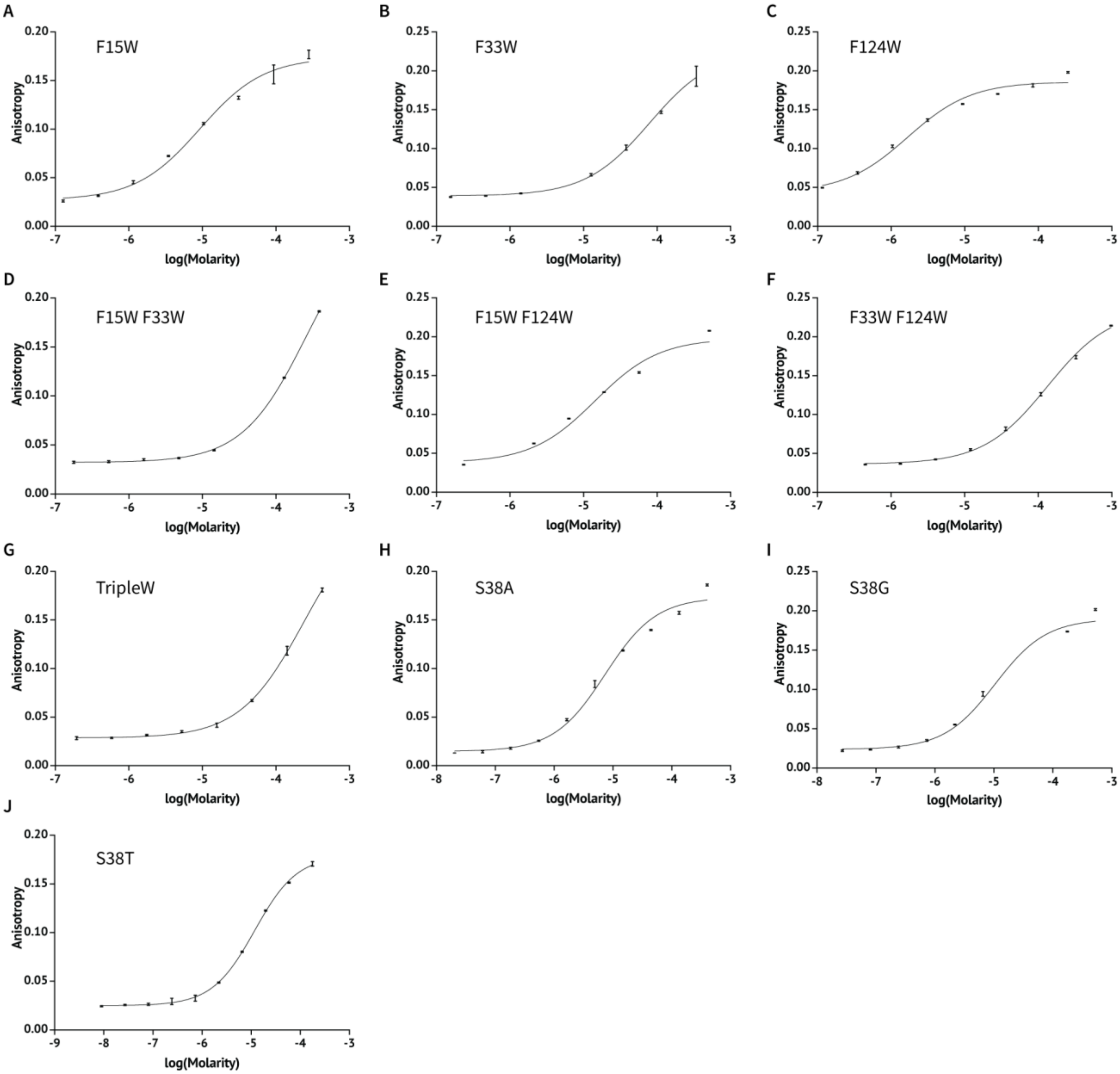


**Supplementary Figure 3. The affinities of mutant G3BP1 NTF2 domains for Caprin1.** Fluorescence anisotropy data were collected using increasing concentrations of G3BP1 NTF2 in the presence of 30 nM reporter peptide derived from the Caprin1 binding motif (*F**-YNFIQDSMLDFE). Data represent the mean and standard deviation of n=3 experiments. For each construct, the average from three independent titrations was fit using a non-linear least-squares algorithm as summarized in Table 2. (A) F15W, (B) F33W, (C) F124W, (D) F15/33W, (E) F15/124W, (F) F33/124W, (G) TripleW, (H) S38A, (I) S38G, and (J) S38T.

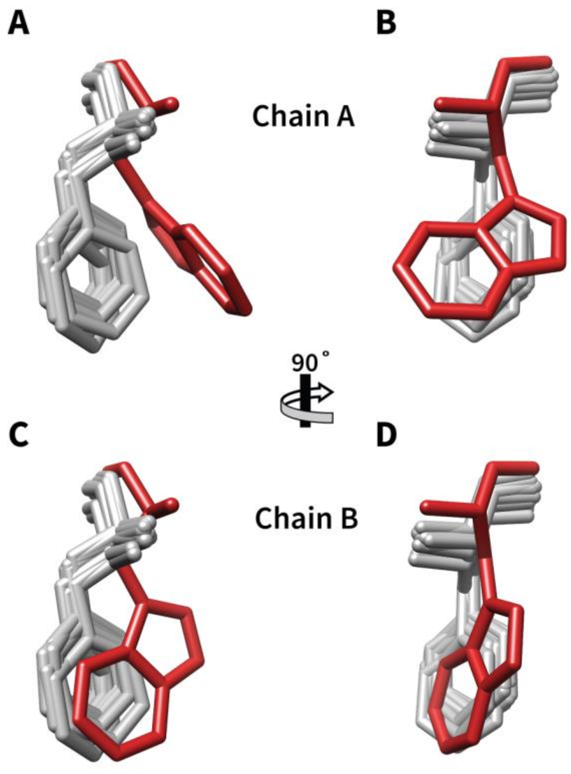


**Supplementary Figure 4. Stereochemical comparison of mutant W33 to native F33.** A cartoon representation of G3BP1 NTF2 residue 33 from Chain A (A & B) and Chain B (C & D). All published G3BP1 NTF2 crystal structures were downloaded and divided into monomers. The G3BP1 NTF2 TripleW was similarly divided into monomers: Chain A and Chain B. All monomers were aligned then every residue was removed, except for residue 33. (A & B) All wild type monomers (grey) overlayed with TripleW Chain A (red). (C & D) All wild type monomers (grey) overlayed with TripleW Chain B (red). PDB IDs: 4FCM, 4FCJ, 6TA7, 5FW5, 3Q90, and 5DRV.

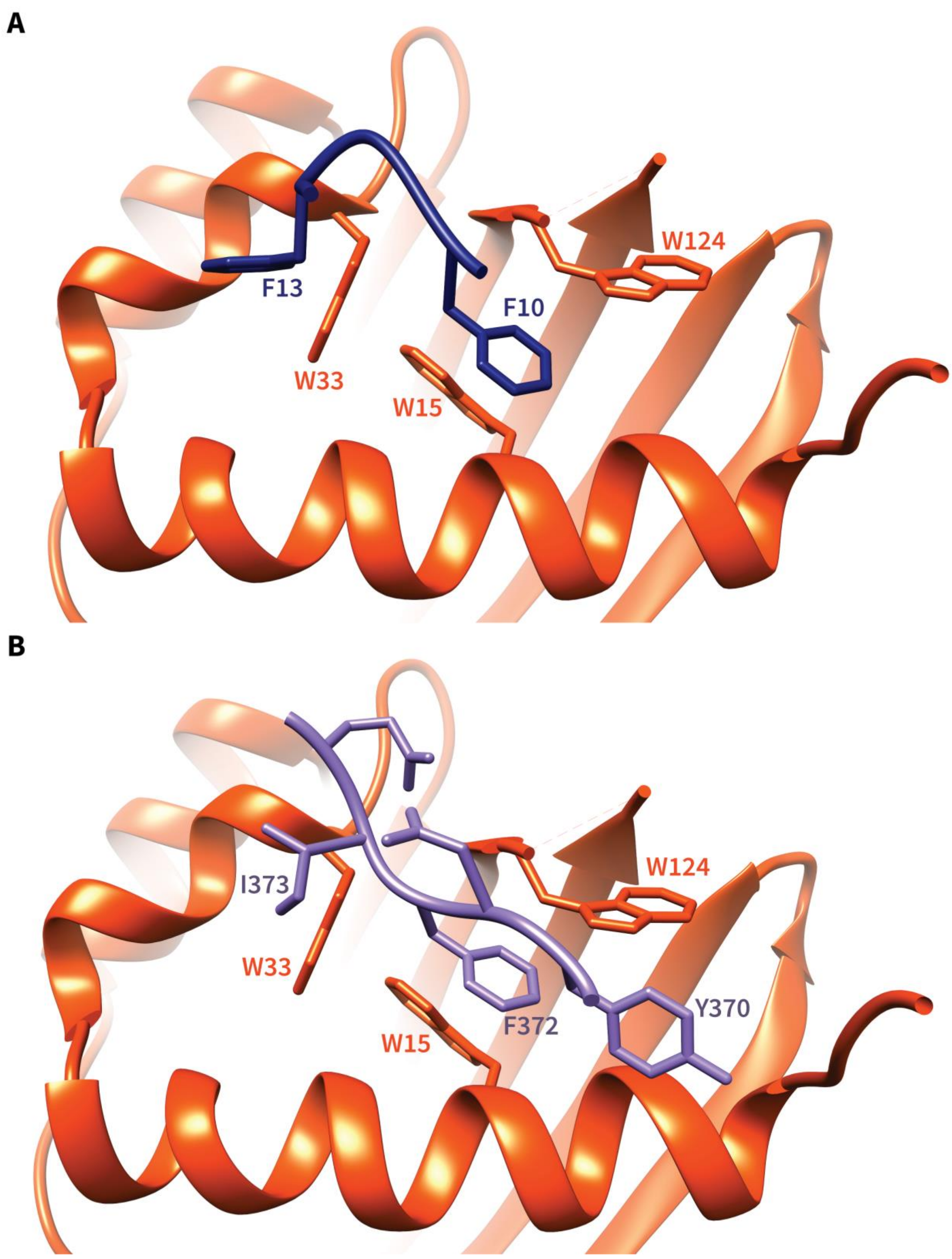
A
F13
F10
W124
W33
W15
B
I373
W124
W33
F372
Y370
W15

**Supplementary Figure 5. Impact of G3BP1 F33W mutation on USP10 and Caprin1 peptide binding in Chain B.** A cartoon representation of G3BP1 NTF2 Triple W Chain B is shown superimposed with the FGDF (A) and YNFIQD peptides (B) from the corresponding WT structures. The NTF2 TripleW structure was aligned with G3BP1 NTF2 WT co-crystal structures, and then the WT NTF2 domain was omitted from the figure. The following co-crystal structures were used: (A) Co-crystal structure of G3BP1 NTF2 WT with USP10 FGDF motif peptide (PDB ID 5DRV), (B) Co-crystal structure of G3BP1 NTF2 WT with Caprin1 YNFIQD motif peptide (PDB ID 6TA7).

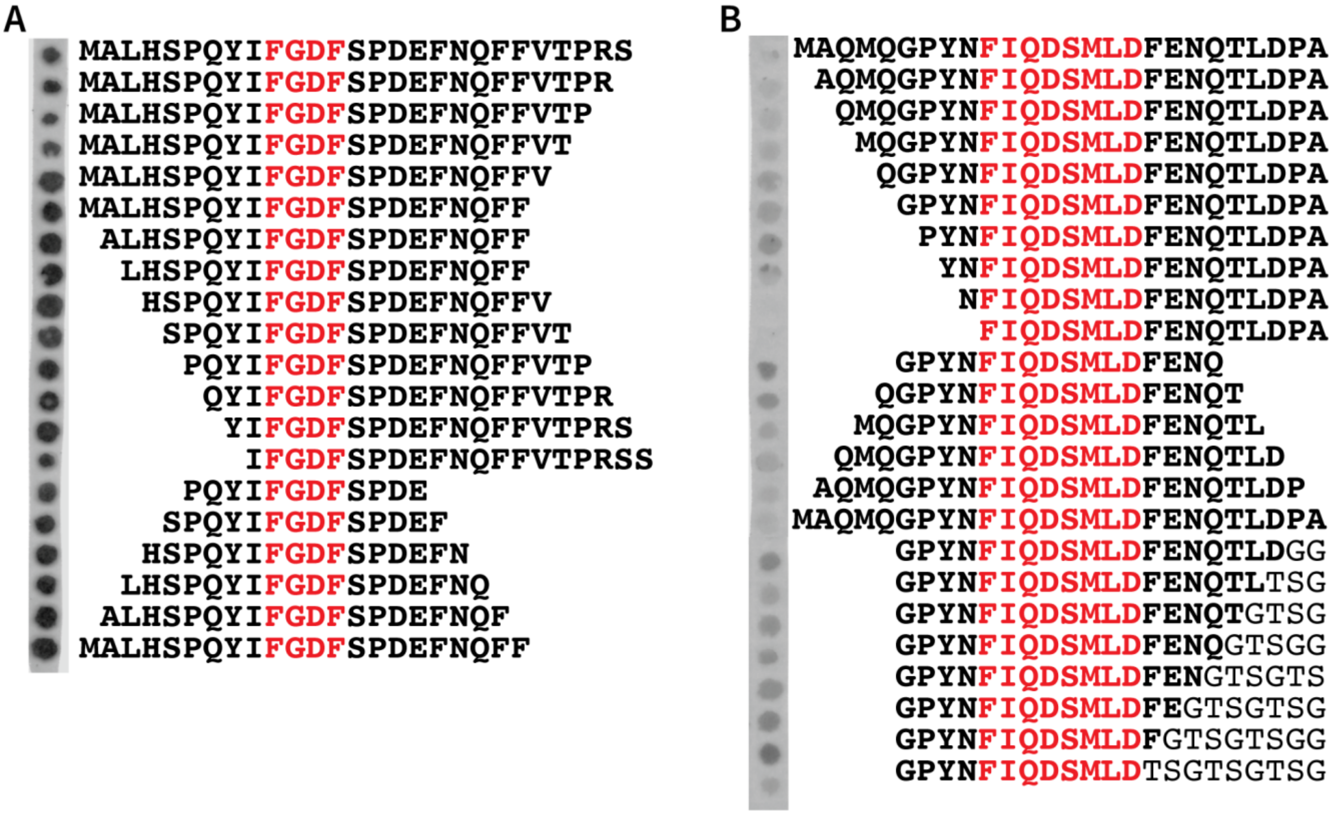


**Supplementary Figure 6. Binding of G3BP1 NTF2 to different length USP10 and Caprin1 peptides.** Peptide cellulose SPOT array with (A) USP10- and (B) Caprin1-derived peptides. Each spot contains the polypeptide listed to the right of the array. Native sequences are in bold and binding motifs are highlighted in red. Darkness represents bound G3PB1 NTF2 protein as detected via antibody staining protocol.

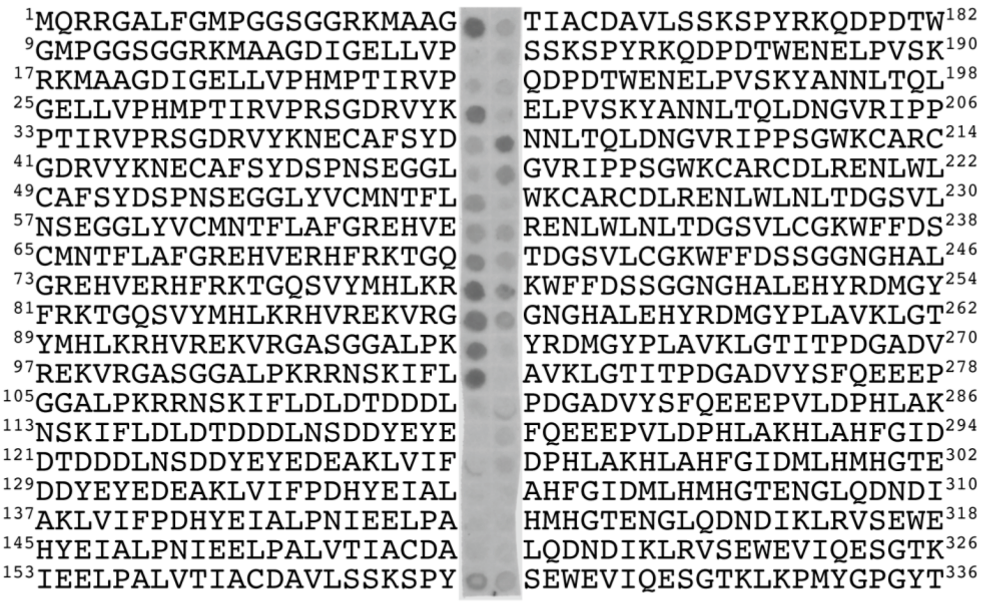


**Supplementary Figure 7. Binding of G3BP1 NTF2 to USP13 amino-acid scan.** Peptide cellulose SPOT array using USP13-derived peptides. Each spot contains the polypeptide listed next to the spot. N- or C-terminal sequence position is labeled for each polypeptide. Darkness represents bound G3PB1 NTF2 protein as detected via antibody staining protocol.

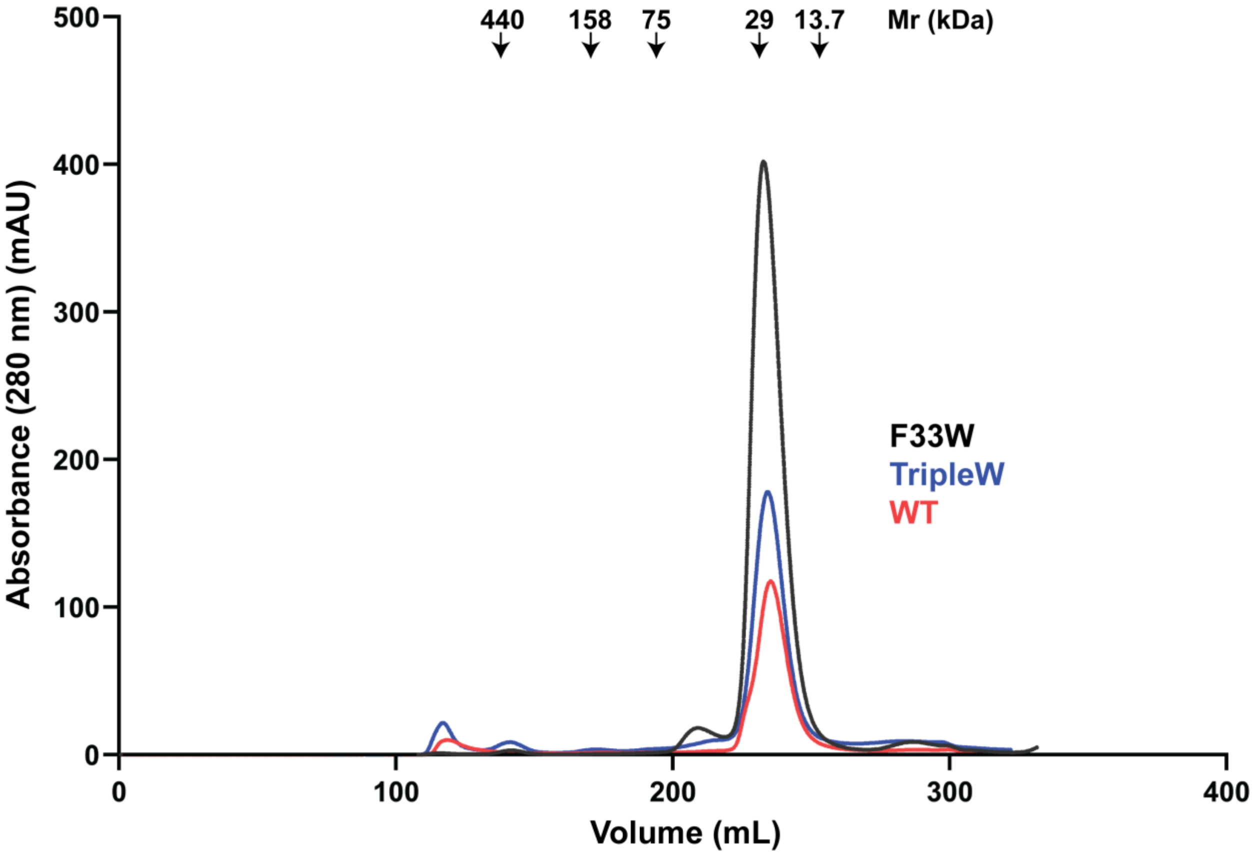


**Supplementary Figure 8. Hydrodynamic homogeneity of recombinantly expressed and purified G3BP1 NTF2-like domain.** Elution chromatograms are shown from a HiLoad 26/600 Superdex 200 pg size-exclusion chromatography column for the WT domain and the F33W and TripleW mutants. Other domains showed similar profiles. The G3BP1 NTF2 protein eluted as a dimer (32 kDa), as calculated using the molecular mass ($M_r$) standards.
